# Intestinal LKB1 loss drives a pre-malignant program along the serrated cancer pathway

**DOI:** 10.1101/2023.07.17.548873

**Authors:** S.F. Plugge, H. Ma, J.Y. van der Vaart, J. Sprangers, F.H.M. Morsink, D. Xanthakis, C. Jamieson, K.B.W. Stroot, A.R. Keijzer, T. Margaritis, T. Candelli, R. Straver, J. de Ridder, F.C.P. Holstege, W.W.J. de Leng, G.J.A. Offerhaus, A. Merenda, M.M. Maurice

## Abstract

**Background & Aims:** Heterozygous inactivating mutations of Serine Threonine Kinase 11 (STK11)/Liver Kinase B1 (LKB1) are causative to the Peutz-Jeghers syndrome (PJS), a hereditary disease characterized by gastrointestinal hamartomatous polyposis and increased cancer susceptibility. While LKB1 loss-induced polyp formation has been ascribed to non-epithelial tissues, how LKB1 deficiency increases cancer risk of patients by altering the phenotypical landscape and hierarchical organization of epithelial tissues remains poorly understood.

**Methods:** Using CRISPR/Cas9, we generated heterozygous and homozygous Lkb1-deficient mouse small intestinal and human colon organoids. These organoids were characterized by an integrated approach that combines imaging, bulk and single-cell RNA sequencing and growth factor dependency assays. Our findings were validated in human PJS-derived tissues using immunohistochemistry and linked to colorectal cancer profiles using the TCGA cancer database.

**Results:** Our results reveal that heterozygous Lkb1 loss is sufficient to push intestinal cells into a premalignant transcriptional program associated with serrated colorectal cancer, which is further amplified by loss-of-heterozygosity. This altered epithelial growth state associates with persistent features of regeneration and enhanced EGFR ligand and receptor expression, conferring niche-independent growth properties to Lkb1-deficient organoids. Moreover, our newly generated LKB1-mutant signature is enriched in sporadic serrated colorectal cancer, and synergistic cooperation of Lkb1-deficiency with mutant Kras was experimentally confirmed by assessing organoid growth properties and transcriptomes.

**Conclusions:** Heterozygous loss of LKB1 pushes intestinal cells into a chronic regenerative state which is amplified upon loss-of-heterozygosity. Lkb1-deficiency thereby generates fertile ground for serrated colorectal cancer formation in the intestine, potentially explaining the increased cancer risk observed in PJS.

## Introduction

Peutz-Jeghers syndrome (PJS) is a hereditary genetic disorder affecting approximately 1:50.000-1:200.000 individuals worldwide^1^. PJS is characterized by the formation of benign hamartomatous polyps throughout the gastrointestinal tract, which necessitates periodical surgical resection. Additionally, patients carry an increased risk of developing early-onset epithelial tumors, particularly within the gastrointestinal tract (up to 66%), breast (up to 54%), pancreas (up to 36%) and gonads^1–3^, resulting in a cumulative lifetime risk of 93%^3^. Moreover, PJS patients are predisposed to small intestinal cancers that rarely develop in the general population (0.03% compared to 13% in PJS)^2,4^.

PJS is caused by heterozygous inactivating mutations in the tumor suppressor gene Liver Kinase B1 (*LKB1/STK11*)^5^. Of note, inactivation of *LKB1* is also linked to the formation of sporadic cancers such as lung carcinoma, melanoma, pancreatic and cervical cancers^6,7^. The *LKB1* gene encodes for a serine/threonine kinase involved in diverse cellular processes, including energy metabolism, cell adhesion, DNA methylation and apicobasal polarity^8–11^. Loss of heterozygosity (LOH) of *LKB1* is a driving force of cancer development, since LOH occurs in over 64% of PJS-derived carcinomas^12,13^.

Recent studies using murine models revealed that *Lkb1* loss in non-epithelial lineages suffices to induce the development of GI polyps resembling those observed in PJS patients^14–17^. This led to a reevaluation of *Lkb1*’s role in non-epithelial tissues, with an emphasis on polyposis formation. The relationship between polyp formation and cancer predisposition in PJS patients however remains highly debated, as LOH and dysplastic transformation of PJS polyps appear to be very rare compared to other polyposis syndromes like familial adenomatous polyposis (FAP)^12,13,18,19^.

Since *Lkb1* loss mediates predisposition to carcinoma development in multiple epithelial tissues irrespective of polyp formation, we hypothesize that epithelial-intrinsic alterations may sensitize these tissues to malignant transformation. Notably, LKB1 deficiency in the intestinal epithelium itself does not mediate polyp formation but was shown to alter secretory cell fate specification^14,20,21^. Despite these advancements, how alterations in intestinal epithelial organization due to *LKB1* predispose for cancer development in PJS patients remains poorly understood.

Here, we model *mono-* and *bi*-allelic inactivation of *Lkb1* in mouse and human intestinal organoids to investigate how *Lkb1* loss affects the intestinal epithelial cellular and transcriptional landscape. We uncover that *Lkb1* loss promotes features of epithelial transformation linked to the serrated pathway for intestinal carcinogenesis. These features include loss of classical adult stem cell populations, activation of regeneration-related transcriptional programs, intestinal-to-gastric cellular transdifferentation, and the acquirement of niche-independent growth properties. Our findings argue that *Lkb1* loss mediates epithelium-wide activation of a pre-cancer program, providing an explanation for the increased cancer risk observed in PJS patients. By linking *Lkb1* loss to a major pathway for colorectal cancer (CRC) development, these findings pave a path for the development of new strategies for prevention, surveillance and precision treatment of PJS tumors and other malignancies harboring *LKB1*-inactivating mutations.

## Materials and Methods

### Patient material

Colorectal tissue collection for the generation of healthy colon organoids was performed according to the guidelines of the European Network of Research Ethics Committees (EUREC). Formalin-Fixed Paraffin-Embedded (FFPE) intestinal tissue samples were obtained from pathology archives. For details see supplementary methods.

### Organoid culture

Both mouse small intestinal organoids and human colon organoids were established and cultured as previously described^22,23^. Organoids were grown in Matrigel (Corning) or BME (Cultrex) and passaged every 6-8 days; their medium was refreshed every 2-3 days. For medium composition see supplementary materials.

### Gene editing of organoids

For generation of mutant and overexpression organoids, electroporation of organoids was performed as previously described^24^. For further details, plasmids and organoid selection see supplementary materials.

### Imaging of organoids

Organoids were fixed and stained 3-4 days after passaging. For detailed protocols on immunofluorescence, immunohistochemistry and electron microscopy see supplementary materials.

### Bulk RNA sequencing

Organoids were lysed in RLT lysis buffer (Qiagen) 3-5 days after passaging. Total RNA was processed for sequencing using Illumina Truseq Stranded mRNA kit and sequenced. For details see supplementary materials.

### Single cell RNA sequencing

Single cells from organoid were harvested using TrypLE, four days after passaging. Cells were filtered and scRNA-seq was performed according to the Sort-seq protocol^25^. For details see supplementary materials.

## Results

### *Lkb1* bi-allelic loss results in phenotypic heterogeneity and increased self-renewal potential of small intestinal organoids

To study the role of *Lkb1* loss in the intestinal epithelium, we employed CRISPR/Cas9 technology to generate *Lkb1*-mutant mouse small intestinal organoid (mSIO) lines. Targeted organoids were cultured transiently in WENR (see supplementary materials) to enhance outgrowth efficiency^26^. Under these conditions, organoids acquire a cystic morphology, due to the expansion of stem cells^22,27^. To obtain genotypically clonal *Lkb1*-mutant cultures, we grew organoids from single cells. Sequencing analysis confirmed the generation of homozygous mutant clones carrying bi-allelic truncating indels (Figure 1A, *Lkb1*^-/-^). In addition, we obtained a heterozygous clone that carried an out-of-frame insertion in one allele and an in-frame deletion (p.P38_R39del) in the other allele. The two deleted amino acids are located outside essential protein regions (Figure 1A, *Lkb1^+/-^*) and allowed for protein expression (Figure 1B), predicting retained protein function. Subsequently, organoids were transferred to ENR medium, allowing for formation of typical budding structures resembling the *in vivo* crypt-villus axis^22^. Clones that transitioned into budding organoids over four passages comprised all genotypes, including wild-type (WT), *Lkb1^+/-^* or *Lkb1^-/-^ (Lkb1^bud-/-^)*. Strikingly, a subset of clones that retained cystic morphology were exclusively genotyped as *Lkb1^-/-^* (*Lkb1^cys-/-^*) (Figure S1A). In line with Lkb1’s role in AMPK-mTOR signaling, we observed reduced phosphorylated Ampk (pAmpk), a direct substrate of Lkb1 kinase activity^10^, along with upregulated mTor signaling, indicated by increased phosphorylated 4E-BP1 in *Lkb1^-/-^* organoids (Figure 1C-D). As expected, *Lkb1^+/-^* organoids displayed an intermediate phenotype, consistent with mono-allelic expression (Figure 1C-D). Furthermore, *Lkb1^bud-/-^*organoid cultures displayed a notable accumulation of dead cells at their basal side (Figure 1E), validated by cleaved caspase-3 staining (Figure 1F). This observation differed from WT and *Lkb1^+/-^* organoids for which dying cells were apically extruded and shed into the lumen, as shown previously (Figure 1F)^22^. Mislocalization of dead cells in *Lkb1^bud-/-^* organoids was corrected by wild-type LKB1 overexpression, indicating that this is a consequence of *Lkb1* loss (Figure S1B-C).

**Figure 1.**
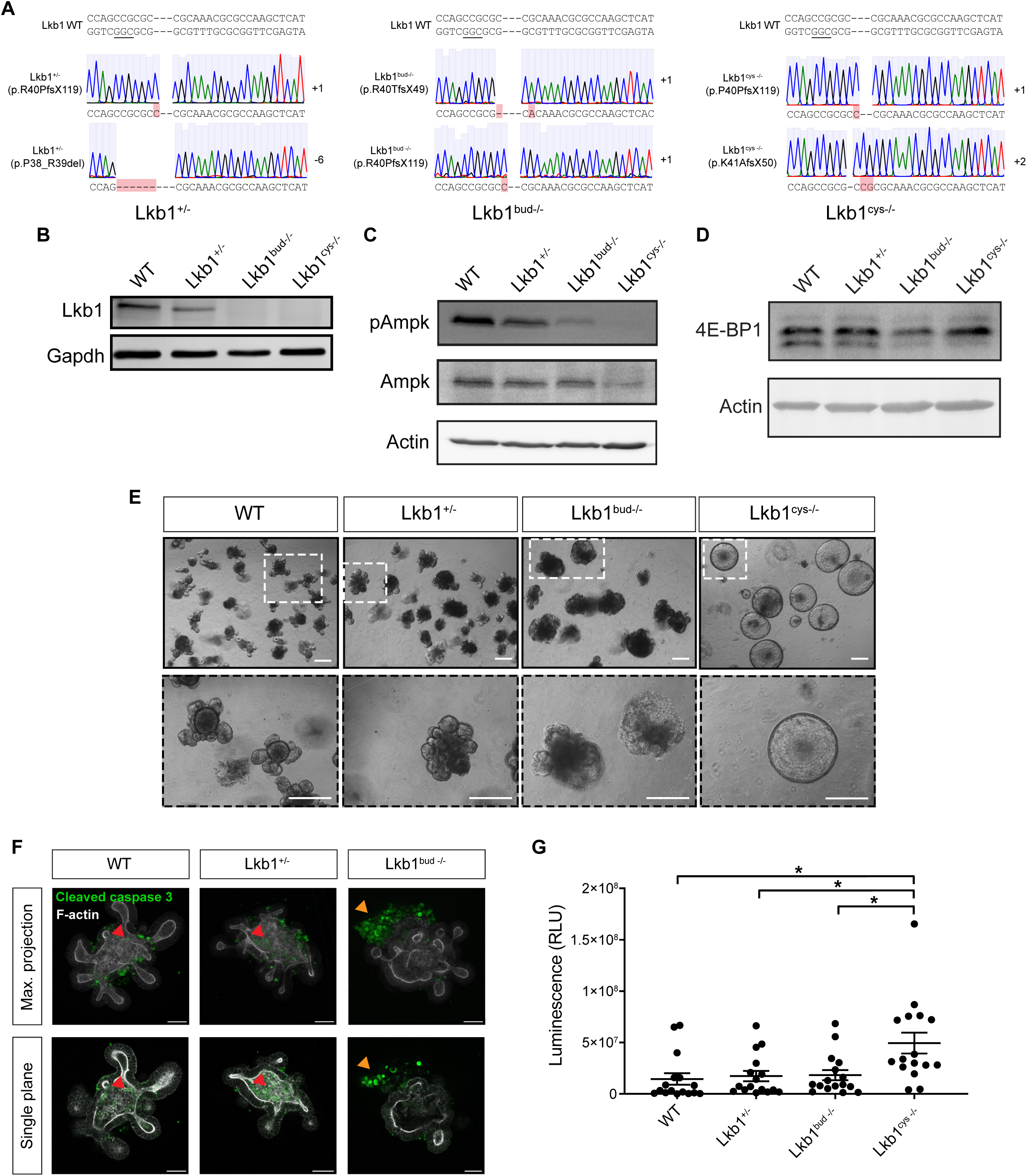
*Lkb1*-mutant organoids display alterations in phenotype and reconstitution capacity. **A)** Sequences of *Lkb1*-mutant organoids obtained with gRNA1 showing indels/deletions detected in exon 1 of *Lkb1*. *Lkb1^+/-^*shows two amino acids deletion outside essential Lkb1 domains. PAM sequences are underlined. **B-D)** Western blot of Lkb1 **(B)**, (p)AMPK **(C)**, and 4E-BP1 **(D)** in WT and *Lkb1*-mutant organoids. **E)** Brightfield images of *Lkb1*-mutant organoid clones. Scale bar = 100 µM. **F)** Immunofluorescent images of cleaved caspase-3 (CC3) and F-actin in *Lkb1*-mutant and WT organoids. Intraluminal (red arrowheads) and extraluminal CC3^+^ cells (orange arrowheads) are indicated. Scale bar = 50 µm. **G)** Organoid reconstitution assay from single cells. One-way ANOVA (Dunnett) was applied. *\** = *p < 0.05*. *N* = 16.

To assess the stability of budding and cystic *Lkb1^-/-^*organoid phenotypes, we generated a *Lkb1^-/-^* bulk population (100% knockout score). One week after transfer to ENR medium, the cystic-to-budding ratio was 1:1 (Figure S1D). We next expanded nine budding and nine cystic clonal organoids lines and tracked their morphology. After two passages, ∼70% of *Lkb1^bud-/-^* clones retained budding morphology, while all *Lkb1^cys-/-^*clones remained cystic (Figure S1E). Thus, homozygous *Lkb1* loss leads to two distinct morphologies, with some budding-to-cystic conversion. Of note, reversal of cystic morphology by reintroduction of LKB1 expression was not possible, as LKB1^+^/mCherry^+^ cells were rapidly lost over time (Figure S1F-G), potentially due to competition by *Lkb1^cys-/-^*cells^28^. Next, we examined whether *Lkb1* loss alters the ability to reconstitute organoids from single cells, as a proxy for their self-renewal capacity. While WT, *Lkb1^+/-^* and *Lkb1^bud-/-^*organoids displayed similar outgrowth rates, *Lkb1^cys-/-^* organoids carried a significantly improved capacity to form organoids (Figure 1G).

To rule out that the observed phenotypes are caused by unintended gRNA-induced mutations rather than *Lkb1* loss, we employed an alternative, non-overlapping *Lkb1*-targeting gRNA (gRNA2). Targeting *Lkb1* with gRNA2 led to similar morphologic changes, including shedding of dead cells at the basal side and a divergent budding and cystic morphology for *Lkb1^-/-^* (Figure S2). Moreover, newly generated *Lkb1^cys-/-^* clones also displayed superior reconstitution potential compared to their *Lkb1^bud-/-^* counterparts (Figure S2C). We conclude that *Lkb1*-deficiency results in specific organoid phenotypes related to loss of one or both alleles of *Lkb1*.

### *Lkb1* loss alters the number, morphology and positioning of secretory cells

To investigate whether our *Lkb1*-mutant organoid lines serve as representative models for intestinal epithelial *Lkb1* loss *in vivo*, we examined alterations in localization and morphology of Paneth (Lyz1^+^) and goblet (Muc2^+^) cells previously reported for the *Lkb1*-mutant mouse intestine^20,21^. Compared to WT organoids, Lyz1^+^ cells in *Lkb1*^+/-^ and *Lkb1^bud-/-^* organoids were scattered beyond the crypt region into the villus domain (Figure 2A, S3A). Additionally, in both WT and *Lkb1^+/-^*organoids, Muc2^+^ cells were sparsely located to the villus region, while *Lkb1^bud-/-^* organoids displayed an increase in Muc2^+^ cells that were found scattered throughout the crypt-villus axis (Figure 2B). These observations resemble reported aberrancies in Paneth and goblet cell numbers and localization in *Lkb1*-deficient mouse intestine *in vivo*^20^. Conversely, *Lkb1^cys-/-^* organoids lacked Lyz1^+^ cells but did form Muc2^+^ cells that are usually lacking in cystic organoids^27^, indicating that secretory lineage specification is also altered within these organoids (Figure 2A-B).

**Figure 2.**
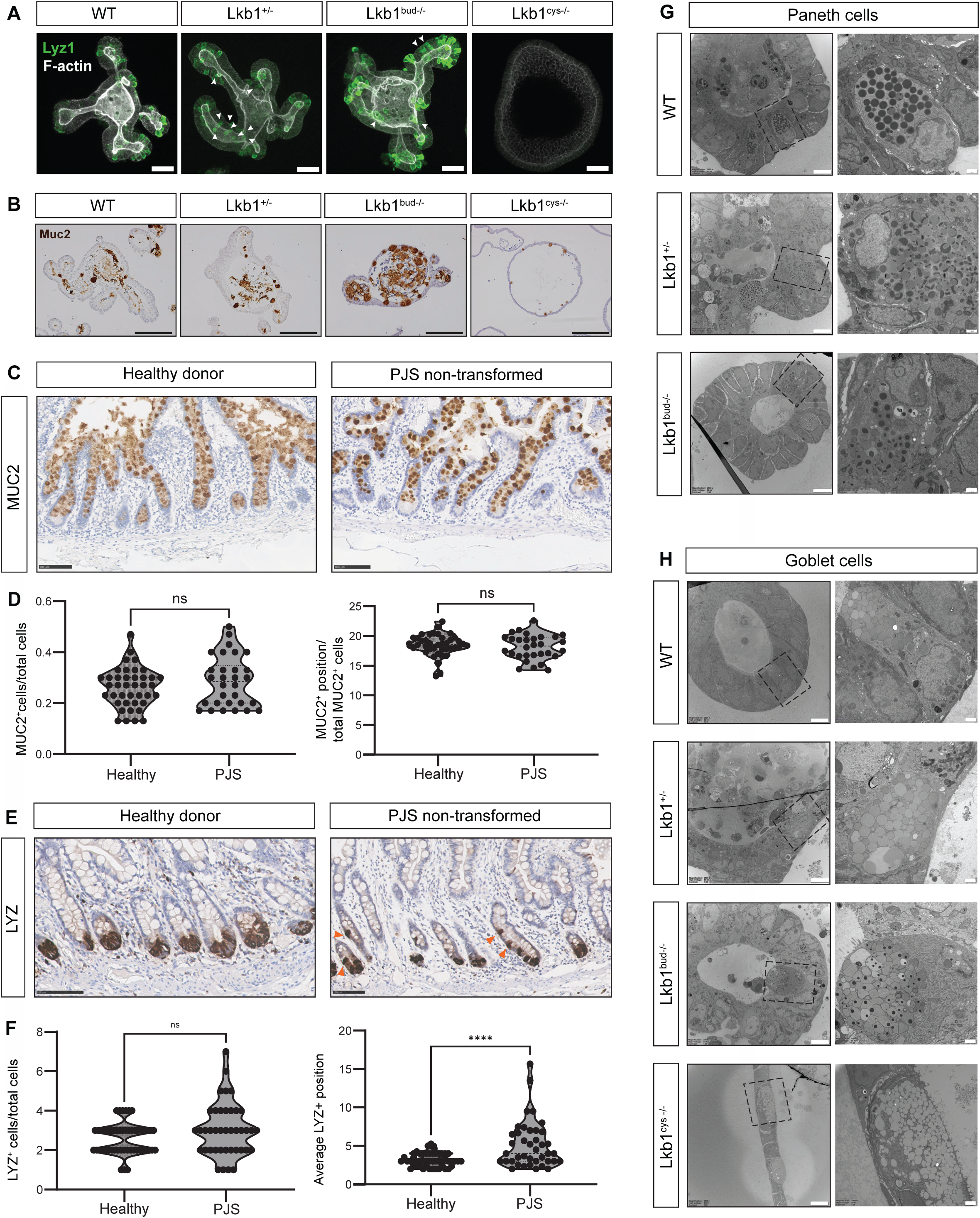
*Lkb1* loss mediates alterations in the number, location and ultrastructural morphology of secretory cells. **A)** Immunofluorescence staining of Lysozyme1 (Lyz1) and F-actin in WT and *Lkb1*-mutant small intestinal organoids. Scale bar = 50 μm. White arrows indicate mislocalized Lyz1^+^ cells. **B)** Immunohistochemistry staining of Mucin 2 (Muc2) in WT and *Lkb1*-mutant organoids. Scale bar = 100 μm. **C&E)** Immunohistochemistry staining of MUC2 **(C)** and LYZ **(E)** in healthy and non-transformed PJS epithelium. Orange arrowheads indicate mislocalized LYZ^+^ cells. Scale bar = 100 μm. Shown images are zoom-in of Figure S3B-C. **D&F)** The average position and number of MUC2+ cells **(D)** and LYZ+ cells **(F)** in healthy and PJS epithelia. Each dot represents a single crypt. Unpaired t-test was applied. *ns = not significant*, *\*\*\*\* = p < 0.0001*. **G-H)** Electron microscopy images Paneth **(G)** and goblet **(H)** cells in WT and *Lkb1*-mutant organoids. Scale bar upper panel = 5 μm, lower panel = 1 μm.

We next compared these findings to the intestinal tissue of human PJS patients. The number and localization of MUC2^+^ cells per intestinal crypt were similar in healthy and PJS patient-derived tissues (Figure 2C-D, S3B). Compared to healthy donors, however, LYZ^+^ cells were mislocalized within the PJS mucosa (Figure 2E-F, S3C). Thus, heterozygous loss of *LKB1* in PJS intestines leads to an altered LYZ^+^ cell distribution, similar to our observations in *Lkb1^+/-^* organoids.

To examine *Lkb1* loss-induced alterations in secretory cell morphology at the ultrastructural level, we employed electron microscopy. In WT intestinal organoids, Paneth cells displayed a typical apical accumulation of spherically-shaped secretory granules filled with electron-dense content (Figure 2G)^22^, while *Lkb1^+/-^* Paneth cells carried granules with an abnormal ‘sausage-shaped’ electron-dense core (Figure 2G). In *Lkb1^bud-/-^* Paneth cell granules, the electron-dense core was surrounded by a peripheral electron-lucent halo (Figure 2G). WT goblet cells were columnar-shaped and contained typical large translucent mucus-filled granules (Figure 2H)^22,29^. *Lkb1^+/-^* mucus-secreting cells however were rounded and flattened while *Lkb1^bud-/-^* goblet-like cells displayed granules with an unusual electron-dense core (Figure 2H). In addition, in *Lkb1^cys-/-^* organoids the mucus-like granule content was frequently lost during EM preparation, suggesting an altered mucin composition and explaining the limited number of observed Muc2^+^ cells (Figure 2H). Morphologically, Paneth- and goblet-like cells in all *Lkb1*-mutant organoids looked like ‘intermediate’ cells that were proposed previously to represent transition states between undifferentiated and mature cells^29^, suggesting that *Lkb1* loss may interfere with terminal differentiation of secretory cells. Furthermore, although previous studies^11^ identified roles of *LKB1* in apical brush border formation using 2D human cell lines, our EM images showed no obvious brush border defects in any of the *Lkb1*-mutant organoids (Figure S3D).

Our findings thus reveal that *Lkb1* loss in budding organoids induces alterations in secretory cell number, localization and morphology similar to *in vivo* mouse models and PJS patient intestines, validating these organoids as tools for modelling epithelial alterations in PJS. While *Lkb1^cys-/-^* organoids are more difficult to directly link to PJS epithelial organization *in vivo*, they may represent an intermediate state in cancer progression. We therefore included all clonal organoid variants for further characterization.

### *Lkb1* mono- and bi-allelic loss induces incremental expression of a regenerative gene signature

To examine how *Lkb1* loss affects the epithelial landscape, we determined transcriptional changes using bulk mRNA sequencing. Compared to WT organoids, we observed differential expression of 1298, 2312 and 7816 genes in *Lkb1*^+/-^, *Lkb1^bud-/-^* and *Lkb1^cys-/-^*organoids, respectively (Table S1).

Using gene set enrichment analysis (GSEA) we examined transcriptional gene profiles of individual intestinal cell types in *Lkb1*-mutant organoids^30^. For clarity’s sake, in the following sections we will use ‘*Lkb1-* mutant’ to refer to all genotypes (both mono-allelic and bi-allelic deficient). All *Lkb1*-mutant clones showed significant downregulation of gene sets linked to mature Paneth cells, whereas goblet cell gene sets were enriched within *Lkb1^bud-/-^* organoids in comparison to WT organoids, again suggesting alterations in secretory cell lineage specification (Figure 3A). Noticeably, the *Lgr5^+^* adult stem cell signature^30^ was downregulated in all *Lkb1*-mutant clones (Figure 3A, S4A), reminiscent of fetal intestinal spheroids that rely on *Lgr5*-negative progenitors^31^. Indeed, we uncovered a gradual enrichment of an embryonic spheroid gene signature^31^ from WT to *Lkb1^+/-^* to *Lkb1^bud-/-^* organoids, correlating with the number of lost *Lkb1* alleles (Figure 3A-B). This transition was further enhanced in *Lkb1^cys-/-^* versus *Lkb1^bud-/-^*organoids (Figure S4A-B). Previous studies linked transient upregulation of a fetal-like intestinal gene signature to epithelial remodeling and collagen deposition during injury repair^32–36^. In line, all *Lkb1*-mutant organoid clones exhibited enrichment of gene signatures associated with intestinal regeneration following damage or infection (Figure 3A, S4A).

**Figure 3.**
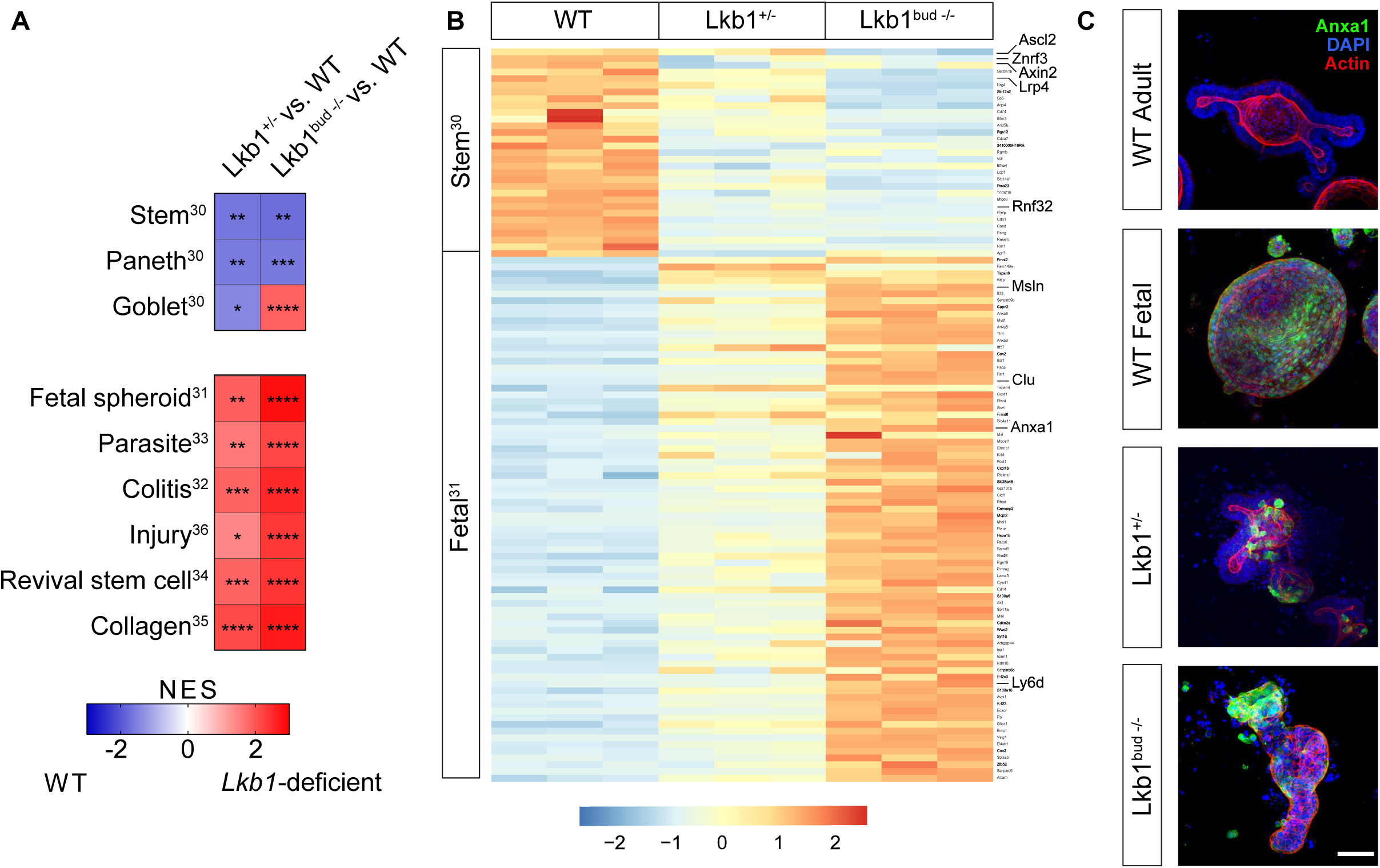
*Lkb1* loss mediates increased expression of a regenerative gene signature in the intestinal epithelium. **A)** GSEA of *Lkb1*-mutant versus WT organoids for cell types^30^, colitis^32^, injury^36^, parasite^33^, fetal spheroid^31^, collagen^35^ and revival stem cell^34^ gene sets. Heatmap displays normalized enrichment scores (NES). ns = not significant, * = FDR < 0.05, ** = FDR ≤ 0.01, *** = FDR ≤ 0.001, **** = FDR ≤ 0.0001. **B)** Heatmap of z-score transformed expression values of *Lgr5^+^* stem cell genes (Stem)^30^ enriched in WT organoids and fetal genes (Fetal)^31^ enriched in *Lkb1*-mutant organoids. Marker genes are highlighted. N = 3. **C)** Immunofluorescence staining of Anxa1, DAPI and F-actin in WT and *Lkb1*-mutant organoids. Fetal organoids from E16.5 mice are shown as a positive control. Scale bar = 50 μm.

To confirm that *Lkb1* deficiency rather than organoid morphology mediates activation of a regenerative program in Lkb1^cys-/-^ organoids, we analyzed transcriptomes of *Apc*^-/-^ and WT Wnt-treated cystic organoids^27^. *Lkb1^cys-/-^*organoids were significantly enriched for expression of regenerative gene sets compared to *Apc^-/-^* and Wnt-treated organoids (Figure S4C). These findings confirm that *Lkb1* loss itself promotes a regenerative state, although morphology may contribute to the enhancement of this phenotype.

Furthermore, all *Lkb1*-mutant clones, and control fetal intestinal organoids (E16.5), displayed increased expression of the fetal marker Anxa1^32^, which was absent in WT adult intestinal organoids (Figure 3C, S4D). These findings indicate that *Lkb1* deficiency converts the epithelium into a primed regenerative state that is already apparent upon monoallelic *Lkb1* loss, which mirrors the PJS intestinal epithelium.

### *Lkb1*-mutant organoids show a Yap-induced regenerative state and display intra-epithelial self-sufficiency for Egf

Over recent years, Yap signaling was placed central to intestinal regeneration^37^. Additionally, *LKB1* was identified as a negatively regulator of the Hippo-Yap pathway^38^. We therefore wondered if *Lkb1* loss may activate Yap signaling in intestinal epithelial organoids. Indeed, we identified enhanced expression of multiple Yap-regulated gene signatures^37,39^ in *Lkb1*-mutant organoids (Figure 4A, S5A). Notably, Yap target gene expression increased with the number of lost *Lkb1* alleles (normalized enrichment scores of 2.01 and 2.68, respectively). Yap target gene expression was further enhanced in *Lkb1^cys-/-^* organoids compared *Lkb1^bud-/-^* organoids (Figure S5A), which is consistent with their enhanced regenerative transcriptional program. Sustained YAP signaling may explain their related cystic morphology (Figure 1E) and Paneth cell deficiency (Figure 2A), as YAP signaling needs to be suppressed to induce symmetry breaking and Paneth cell formation in mSIOs (Figure S5A)^40^. Immunofluorescence staining confirmed increased Yap protein expression and nuclear localization in both *Lkb1^bud-/-^* and *Lkb1^cys-/-^*organoids (Figure 4B-C, S5B-D).

**Figure 4.**
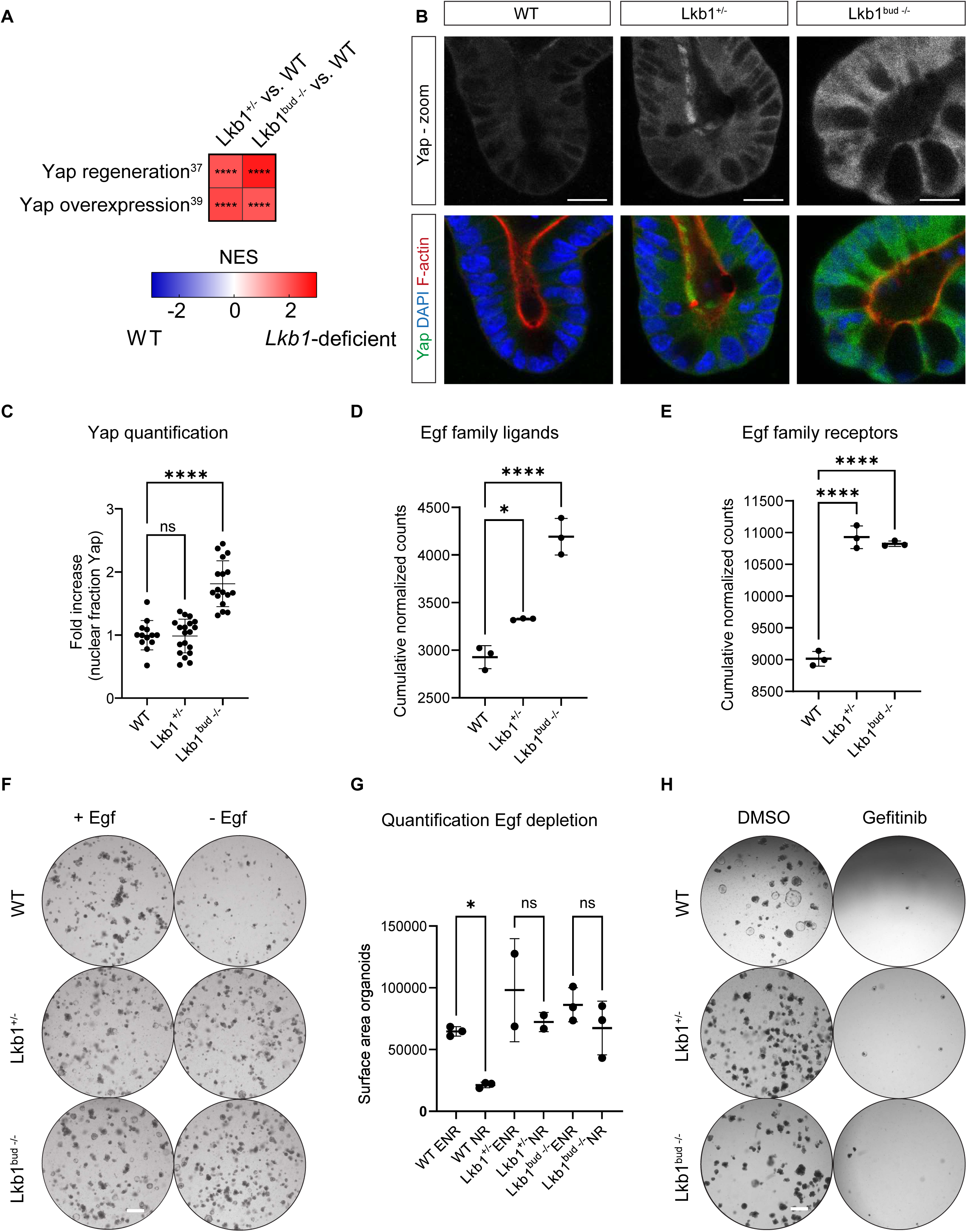
*Lkb1* loss accommodates Yap pathway activation and mediates upregulation of EGF family receptors and their ligands to drive niche independency. **A)** GSEA of *Lkb1*-mutant versus WT organoids for Yap gene sets^37.39^. Heatmap displays normalized enrichment scores (NES). ns = not significant, * = FDR < 0.05, *** = FDR ≤ 0.001, **** = FDR ≤ 0.0001. **B)** Immunofluorescence staining for Yap, DAPI and F-actin in WT and *Lkb1*-mutant organoids. Scale bar = 172 µm. **C)** Quantification of the nuclear fraction of Yap. *N* = 13-19. One-way ANOVA was applied. **D-E)** Cumulative normalized counts of *Egfr* ligands **(D)** and *Egf* family receptors **(E)** in WT and *Lkb1*-mutant organoids. One-way ANOVA was applied. *N* = 3. **F)** Brightfield images of WT and *Lkb1*-mutant organoids grown in the presence (ENR) and absence (NR) of Egf. Scale bar = 500 µm. **G)** Surface area covered by WT and *Lkb1*-mutant organoids grown in ENR or NR. *N* = 2-3. Two-way ANOVA was applied. **H)** Brightfield images of WT and *Lkb1*-mutant organoids cultured in NR and treated with DMSO or 100 nM Gefitinib. Scale bar = 500 µm. *\* = p < 0.05,* and *\*\*\*\* = p < 0.0001*.

YAP pathway activation is well-known to enhance EGF family ligand expression during regeneration^37,41^. In line, *Lkb1*-mutant organoids showed upregulated expression of Egf ligands and receptor family members, correlating with *Lkb1* allelic loss and alterations in organoid morphology (Figure 4D-E, S5E-F). Given the dependency of healthy intestinal organoids on Egf, we examined if *Lkb1*-mutant organoids may grow without Egf. Unlike WT organoids, all *Lkb1*-mutant organoids grew out without Egf supplementation (Figure 4F-G, S5G-H). To ensure this was due to endogenous ligand secretion and not downstream pathway activation, we treated the organoids with Gefitinib, an EGFR inhibitor. All organoid lines remained sensitive to EGFR inhibition (Figure 4H, S5I), confirming endogenous EGF production as a driver of growth. Interestingly, increased Egf ligand expression was linked previously to enhanced secretory cell differentiation^42,43^. Indeed, increased secretory cell differentiation in *Lkb1*-mutant organoids (Figure 2) correlates with increased Egf ligand expression (Figure 4D).

These results demonstrate that *Lkb1* loss induces a Yap-induced regenerative state leading to EGF ligand self-sufficiency. As niche-independent growth is a hallmark of cancer progression^44^, these results suggest that *Lkb1*-mutant epithelial cells carry a growth advantage which may promote tumorigenesis initiation.

### *Lkb1*-deficiency mediates reprogramming of intestinal stem cells into a regenerative state

To analyze how *Lkb1* loss alters intestinal lineage specification and cellular hierarchies, we subjected WT and *Lkb1*-mutant organoids to single cell RNA sequencing. We included mouse fetal intestinal organoids, to explore a potential overlap with fetal gene expression. Unsupervised clustering identified 19 distinct clusters (Figure S6A). *Lkb1^cys-/-^*organoids and fetal cells clearly separated, while *Lkb1^bud-/-^*and *Lkb1^+/-^* organoids more closely resembled WT cells (Figure 5A). These findings indicate that although *Lkb1^cys-/-^* organoids express a fetal-like gene program they do not fully convert into a fetal state (Figure 5A). Furthermore, *Lkb1^cys-/-^* organoids harbor a homogeneous cell population distinct from budding organoid clusters, lacking a known intestinal cellular identity. We therefore annotated cell types only in budding organoid lines, using intestinal cell type gene signatures (Figure 5B, S6B-I)^30^. As the gene signatures for Paneth and goblet cells mapped to the same cluster (Figure S6E, S6H), we marked this cluster as secretory progenitors, similar to previous reports^45^. We reclustered all cells from budding organoids (WT, *Lkb1^+/-^* and *Lkb1^bud-/-^*) for further analyses (Figure 5C-D). Cell type fraction analysis revealed an increased fraction of secretory progenitor cells in *Lkb1^bud-/-^*organoids (Figure 5E), confirming alterations in secretory cell differentiation. This increase corresponded with a decrease in enterocyte numbers, suggesting a shift in differentiation from absorptive towards secretory lineages (Figure 5E). Furthermore, we identified ‘intermediate’ cells in *Lkb1^+/-^* and *Lkb1^bud-/-^* organoids that co-express features of Paneth and goblet cells (Figure 5F), in line with our ultrastructural morphological analysis.

**Figure 5.**
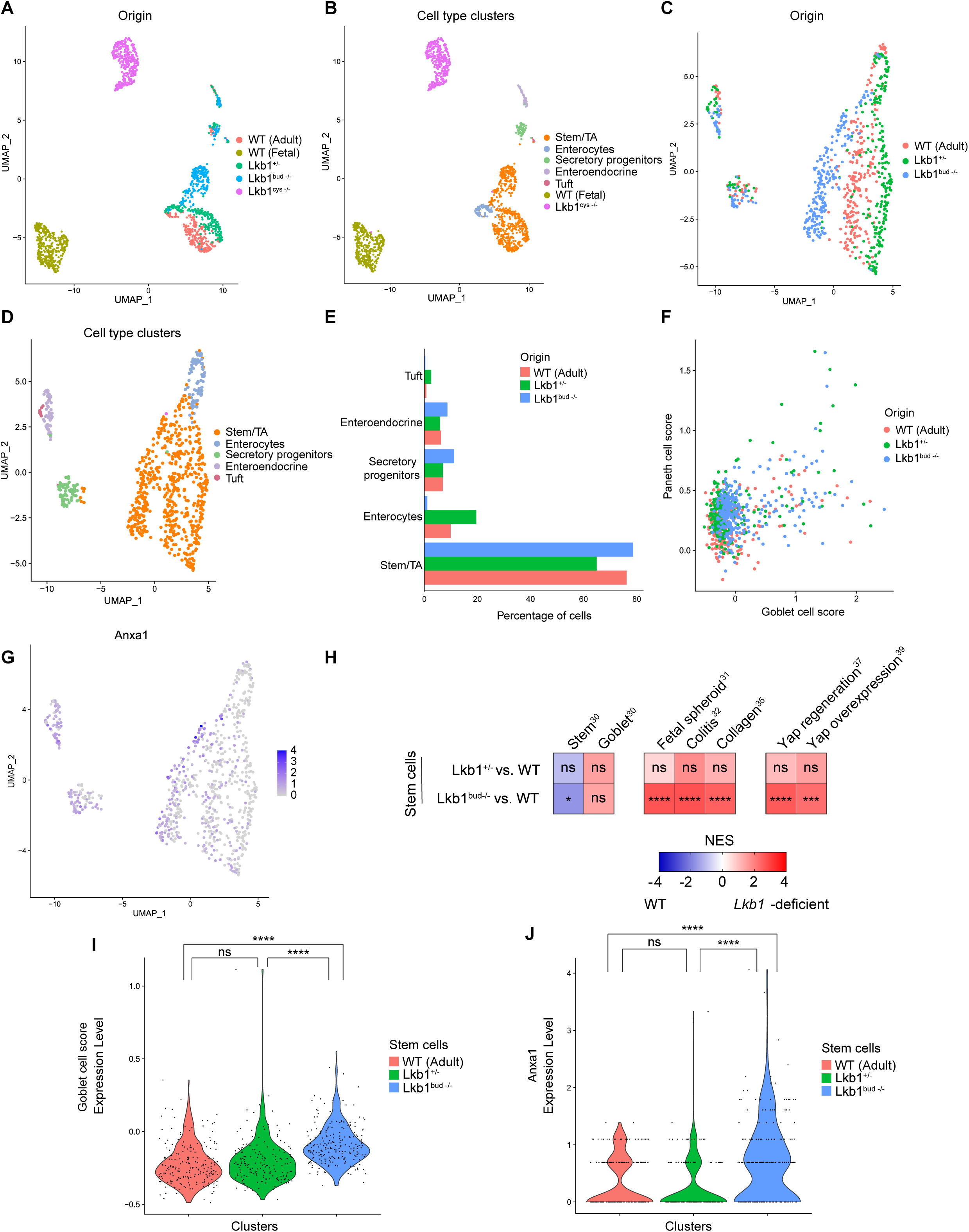
*Lkb1* loss modifies intestinal stem cell identity. **A-B)** UMAP of scRNA-seq data from WT Adult, WT fetal and *Lkb1*-mutant organoids. Colors indicate the different organoids lines **(A)** and the different cell types **(B)**. **C-D)** UMAP of scRNA-seq data from reclustered WT Adult, *Lkb1^+/-^*and *Lkb1^bud-/-^* organoids. Colors indicate the different organoids lines **(C)** and the different cell types **(D)**. **E)** Percentage of cell types for each organoid line. **F)** Scatter plot showing module scores for Paneth^30^ and goblet^30^ cell signatures across WT Adult, *Lkb1^+/-^* and *Lkb1^bud-/-^* cells. Color indicates the origin of the cell. **G)** Expression of *Anxa1* onto the UMAP plot. **H)** GSEA on genes differentially expressed between WT and *Lkb1*-mutant stem cells for cell types^30^, colitis^32^, embryonic spheroid^31^, Yap regeneration^37^, Yap overexpression^39^ and revival stem cell^34^ gene sets. Heatmap displays normalized enrichment scores (NES). ns = not significant, * = FDR < 0.05, *** = FDR ≤ 0.001, **** = FDR ≤ 0.0001. **I-J)** Violin plot showing module scores for goblet cell gene signature^30^ **(I)** or *Anxa1* **(J)** across WT, *Lkb1^+^*^/-^ and *Lkb1^bud-/-^* stem cells. ns = not significant, **** = p ≤ 0.0001 using one-way ANOVA with Wilcoxon test.

Notably, *Anxa1* expression in *Lkb1^bud-/-^* organoids was mainly enhanced within stem and transit amplifying (stem/TA) cell clusters, indicating that these progenitor populations drive the epithelium’s regenerative state (Figure 5G). Reclustering of stem/TA cell populations showed distinct clusters for WT, *Lkb1^+/-^* and *Lkb1^bud-/-^* stem/TA cells (Figure S6J-K). GSEA on genes differentially expressed between WT stem cells and *Lkb1^+/-^ and Lkb1^bud-/-^* stem cells (Table S2) revealed enrichment of goblet cell gene expression within *Lkb1^+/-^ and Lkb1^bud-/-^* stem cell populations (Figure 5H-I). Furthermore, *Lkb1^+/-^ and Lkb1^bud-/-^* stem cells displayed enrichment of regenerative and Yap gene signatures, depletion of the *Lgr5^+^* stem cell signature, and increased expression of Anxa1 (Figure 5H, J). These findings support a model in which *Lkb1* loss induces a regenerative response originating from the stem cell compartment.

In conclusion, our results argue that *Lkb1*-mutant stem/TA cells reside in a regenerative state and preferentially differentiate towards secretory cell lineages.

### *Lkb1* loss drives a pre-malignant program along the serrated pathway which is amplified by additional *Kras* mutations

Next, we aimed to obtain insight in how transcriptional reprogramming induced by *Lkb1* loss may promote cancer predisposition. CRC mainly develops via two pathways: the classical adenoma-carcinoma sequence, initiated by bi-allelic *APC* mutations, and the serrated cancer pathway, initiated by *BRAF* or *KRAS* mutations^46,47^. We examined expression of gene sets linked to these CRC routes to assess if *Lkb1* loss drives entry into one of these pathways. *Lkb1*-mutant organoids displayed significant enrichment of genes that mark neoplastic populations of serrated-specific cells (SSC^46^, iCMS3^48^), which increased with the number of lost *Lkb1* alleles (normalized enrichment scores of 1.70 and 2.16, respectively) (Figure 6A, S7A). Single cell analysis confirmed this trend, attributing the increase of the serrated gene program to *Lkb1^+/-^ and Lkb1^bud-/-^* stem cell populations (Figure S8). This increase was accompanied by the loss of adenoma-specific genes (ASC^46^, iCMS2^48^) (Figure 6A, S7A) and an upregulation of expression of gastric markers like *Aqp5*, *Tacstd2* [*Trop2*], *Tff2*, and *Msln* (Figure 6B, S7B, Table S1), indicating the occurrence of intestinal-to-gastric cellular transdifferentiation^46^. Notably, *LKB1* expression levels were significantly decreased within sessile serrated lesions (SSLs)^46^ compared to pre-cancerous conventional adenomas (AD)^46^ (Figure 6C), supporting the observation that a decrease in LKB1 function accommodates SSL formation.

**Figure 6.**
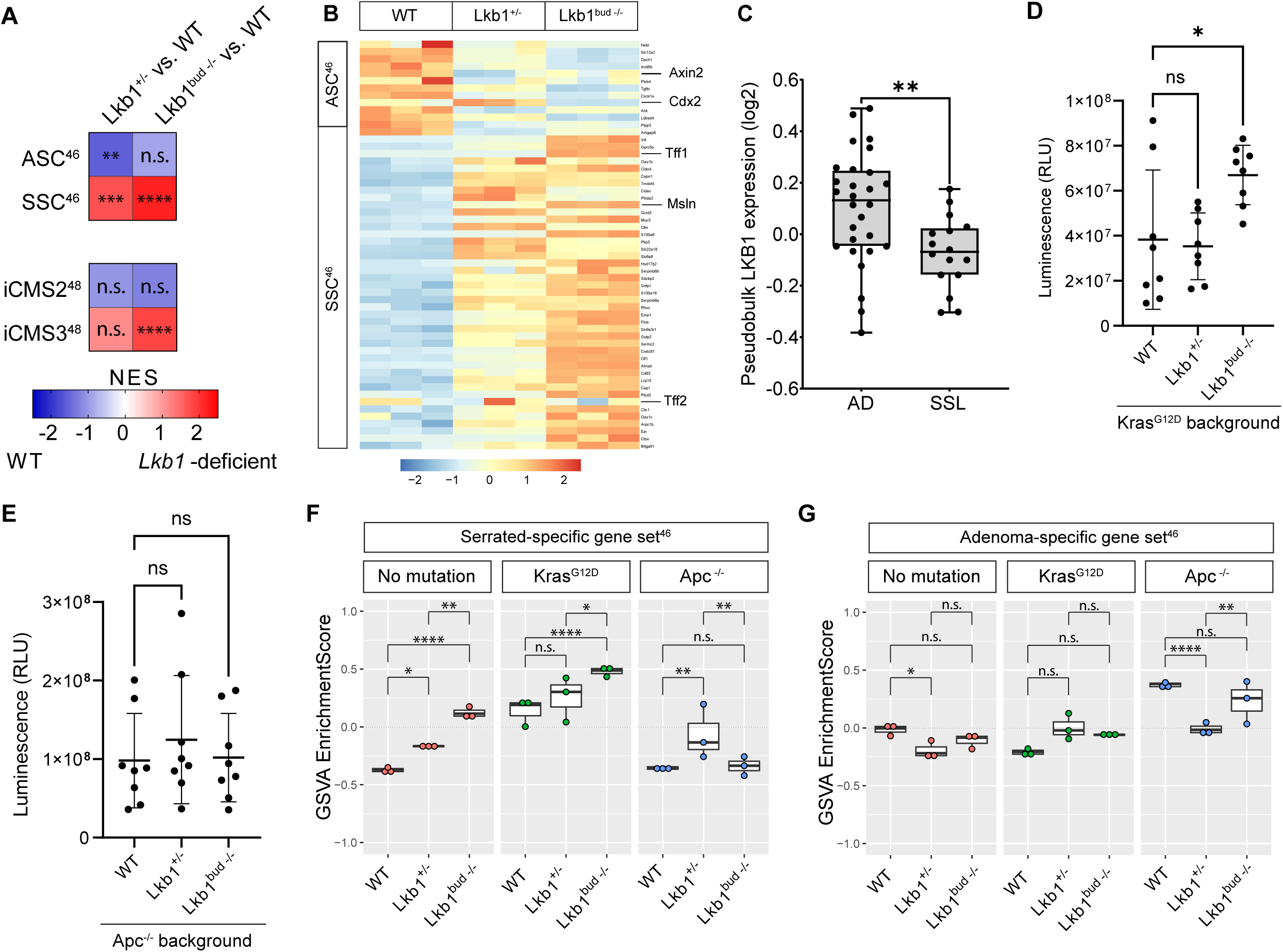
*Lkb1* loss drives a pre-malignant program along the serrated pathway that is further amplified by *Kras* mutations. **A)** GSEA of *Lkb1*-mutant versus WT organoids for serrated-specific (SSC)^46^, adenoma-specific (ASC)^46^, iCMS2^48^ and iCMS3^48^ genesets. Heatmap displays normalized enrichment scores (NES). ns = not significant, ** = FDR < 0.01, *** = FDR ≤ 0.001, **** = FDR ≤ 0.0001. **B)** Heatmap of z-score transformed expression values of ASC^46^ genes enriched in WT organoids and SSC^46^ genes enriched in *Lkb1*-mutant organoids. N = 3. **C)** Boxplot showing log2 transformed *LKB1* expression in pseudobulk samples of pre-malignant adenoma (AD) and sessile serrated lesions (SSL)^46^. **D-E)** Organoid reconstitution assay from single cells for *Kras*-mutant and *Kras*-mutant/*Lkb1*-mutant organoids **(D)** or *Apc*-deficient and *Apc/Lkb1*-mutant organoids **(E)**. *N* = 8 **F-G)** Gene Set Variation Analysis (GSVA) analysis of SSC^46^ genes **(F)** and ASC^46^ genes **(G)** for WT, *Apc* mutant, *Kras* mutant, *Lkb1*-mutant, *Apc/Lkb1* double mutant and *Kras*/*Lkb1* double mutant organoids. *N* = 3. One-way ANOVA with Wilcoxon test was applied. *ns = not significant, * = p < 0.05, ** = p < 0.01* and *\*\*\*\* = p < 0.0001*.

To investigate the susceptibility of an *Lkb1*-mutant epithelial state to early driver mutations found in different CRC subtypes, we introduced *Apc* (Figure S7C) or *Kras* mutations (Figure S7D) in *Lkb1*-mutant organoids. *Kras* mutations synergistically enhanced *Lkb1^bud-/-^* organoid formation from single cells, while *Apc* mutations did not (Figure 6D-E, S7E-F). Bulk mRNA sequencing of double-mutant organoids showed that *Lkb1* loss acts synergistically with Kras^G12D^ mutations to drive a stepwise SSC transcriptomic shift that correlated with allelic loss (Table S3, Figure 6F, S7G). By contrast, no synergism of *Lkb1* loss with ASC signature expression or with *Apc^-/-^* mutations was observed (Figure 6G, S7H).

These results indicate that *Lkb1* loss drives a state of cancer predisposition along the serrated CRC pathway, which is amplified by *Kras* mutations in *Lkb1^+/-^* and *Lkb1^bud-/-^* organoids. Furthermore, *Lkb1^cys-/-^* organoids achieve an advanced serrated state without additional oncogenic mutations. Thus, loss of *LKB1* primes the intestinal epithelium for progression along the serrated pathway.

### LKB1-mutant phenotypes are conserved in human colon organoids and correlate with serrated features in sporadic CRC

To examine if the Lkb1-mutant phenotypes observed in mSIOs can be translated to the human colon epithelium, we obtained one *LKB1^+/-^* and two *LKB1^-/-^*clonal human colon organoid (HCO) lines for which we confirmed a decrease in LKB1 protein expression. In line, pAMPK levels were decreased in *LKB1^-/-^*, but higher in *LKB1^+/-^* HCOs, indicating retained function of the in-frame edited *LKB1* allele (Figure 7A-D). Furthermore, progressive loss of *LKB1* induced enrichment of secretory gene signatures^49^ and multiple regenerative gene sets^31–34,37,39^, similar to our observations in mSIOs (Figure 7E, Table S4). *LKB1* loss also induced EGF self-sufficiency in HCOs, indicated by an increased expression of EGF receptor family members (in both *LKB1^+/-^*and *LKB1^-/-^*; Figure 7F) and EGF ligands (in *LKB1^-/-^*; Figure 7G) as well as an increased tolerance for EGF withdrawal (Figure 7H-I). Notably, all *LKB1*-mutant HCOs retained sensitivity to Gefitinib treatment (Figure 7I). Finally, *LKB1*-mutant HCOs also displayed enrichment of gene signatures linked to the serrated pathway (Figure 7J).

**Figure 7.**
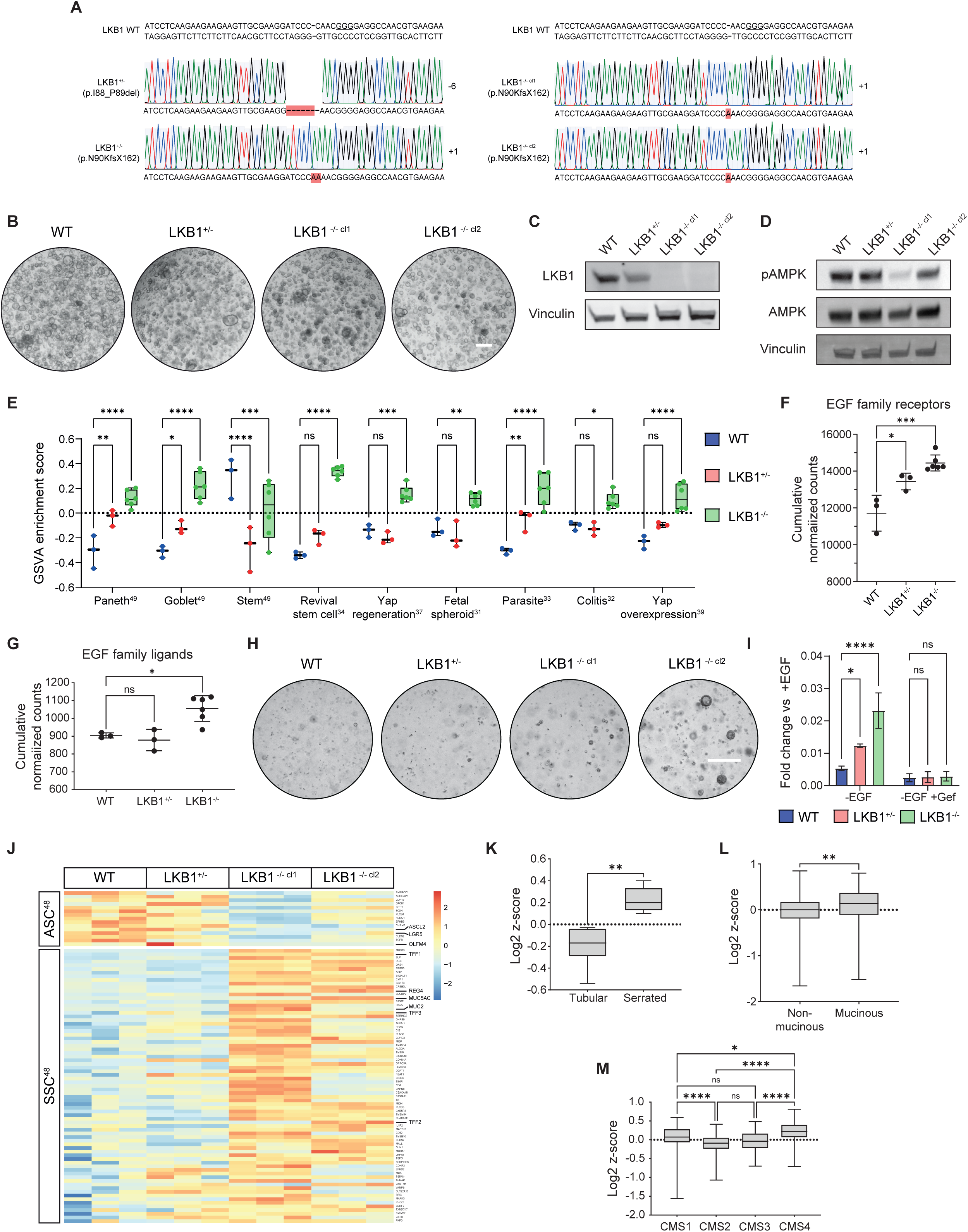
*LKB1* loss phenotypes are conserved in human colon epithelium and sporadic CRC. **A)** CRISPR/Cas9-induced sequence alterations in exon 1 of the *LKB1* gene in human colon organoids (HCOs). *LKB1^+/-^* HCOs acquired a two amino acid deletion in one allele, located outside known functional LKB1 domains. PAM sequences are underlined. **B)** Brightfield images of WT and LKB1-mutant HCOs. Scale bar = 500 µm. **C-D)** Western blot of LKB1 **(C)** and (p)AMPK **(D)** in WT and *LKB1*-mutant HCOs. **E)** Gene Set Variation Analysis (GSVA) analysis of signatures^31–34,37,39,49^ enriched in *LKB1*-mutant HCOs. *N* = 3-6. One-way ANOVA with Wilcoxon test was applied. *ns = not significant, ** = p < 0.05, ** = p < 0.01, *** = p < 0.001* and *\*\*\*\* = p < 0.0001.* **F-G)** Cumulative normalized counts of EGF receptors **(F)** and EGF ligands **(G)** in WT and *LKB1*-mutant HCOs. One-way ANOVA was applied. *N* = 3-6. **H)** Brightfield images of WT and *LKB1*-mutant HCOs grown in the absence of EGF. Scale bar = 500 µm. **I)** Single cell outgrowth assay with and without EGF. One-way ANOVA (Dunnett) was applied. *N* = 3. **J)** Heatmap of z-score transformed expression values of ASC^46^ genes enriched in WT organoids and SSC^46^ genes enriched in *LKB1*^-/-^ organoids. *N* = 3-6. **K-M)** Box plots showing log2 Z-scores for the *LKB1*-mutant signature over the bulk RNA sequencing datasets from Sousa E Melo *et al*^50^. **(K)** or TCGA **(L-M)** grouped as tubular (*N* = 7) or serrated (*N* = 6) **(K)**, mucinous (*N* = 68) or non-mucinous (*N* = 437) **(L)** or consensus molecular subtype (CMS1; *N* = 63, CMS2; *N* = 125, CMS3; *N* = 45, CMS 4; *N* = 73) **(M)** Welch’s unequal variances t-test or one-way ANOVA with Wilcoxon test was applied. *ns = not significant, * = p < 0.05, ** = p < 0.01 and **** = p < 0.0001*.

To assess how these findings correlate with human CRC development, we derived an *LKB1*-mutant gene signature from our HCO models using a directional strategy, taking gene dosage as a covariate, and assessed its correlation with human CRC datasets. Expression of the *LKB1*-mutant gene signature was found enriched within human SSLs compared to tubular adenomas^50^ (Figure 7K), within mucinous CRC (Figure 7L), a subtype linked to SSLs^51^, and in serrated consensus molecular subtypes CMS1 and CMS4 (Figure 7M)^47^.

Together, these data underscore the relevance of our organoid-derived models. Our findings show that *LKB1* loss-induced phenotypes are conserved among species in the gastrointestinal tract and display shared identity with SSLs in human patients.

## Discussion

Over recent years, *Lkb1*-deficient mouse models have faithfully mimicked intestinal hamartomatous polyposis, a hallmark of PJS disease, suggesting that non-epithelial loss of *LKB1* function drives PJS polyposis^14–17^. The relationship between polyposis and epithelial cancer predisposition in PJS however remains highly debated and does not explain extra-intestinal cancer development^2,12,13,18^. Furthermore, the impact of heterozygous and homozygous *Lkb1* loss on epithelial tissue organization at the single cell level has not been addressed.

We employed mSIOs and HCOs to understand how *Lkb1* loss affects epithelial cell identities and drives a pre-malignant state for CRC. By generating heterozygous and homozygous *Lkb1* knockouts, we aimed to model cancer predisposition and development in PJS patients, as over 64% of PJS-derived carcinomas present inactivation of the second gene copy^12,13^. Our findings argue that heterozygous *Lkb1* loss is sufficient to push cells into a premalignant program along the serrated CRC pathway, independent of the micro-environment. This premalignant program is defined by a sustained regenerative response, niche-independent growth, and aberrant secretory cell differentiation, which is further amplified upon loss of the second allele. If this epithelial-intrinsic phenotypic shift can be fully attributed to the known functions of LKB1^10,38^, or whether it is further amplified by LKB1^+/-^ mesenchyme-induced polyposis in PJS patients requires further investigation.

Our organoid-based model validates the link between *Lkb1* loss and aberrant secretory cell specification *in vivo*. We observed that *Lkb1*-deficiency mediates an increase in goblet cells and mucus formation and the generation of secretory cells with ‘intermediate’ features, both morphologically and transcriptionally. Supporting this, our newly generated *LKB1*-mutant gene signature correlated with transcriptional profiles of mucinous subtypes of sporadic CRC. Similar secretory cell alterations induced by *Lkb1* loss may occur in other epithelial tissues, as mucinous carcinomas are common in PJS patients^52,53^. Although the underlying mechanism requires further investigation, these findings indicate a link between secretory cell alterations and cancer subtype progression.

Transcriptomic analysis revealed that *Lkb1* loss induces a dosage-dependent expression of a regenerative epithelial program. This tissue repair program was activated in standard culture conditions, without external damage, suggesting an inherent cellular response upon *Lkb1* loss. We anticipate that, in PJS patients, external injury events that inflict on the *LKB1*-mutant intestine may enhance this response and lock the epithelium into a chronic regenerative state, which may contribute to increased tumor development. The regenerative state also showed increased *Egf* receptor family and ligand expression, endowing niche-independent properties, which is regarded as a hallmark of cancer^44^. The activation of regenerative programs has been extensively linked to an increased risk of tumorigenesis in many epithelial organs^54–56^, including pancreas and breast. Thus, induction of a chronic regenerative state by *Lkb1* loss may potentially explain the increased risk of cancer development in other organs in PJS patients as well.

Importantly, we uncovered that *Lkb1* loss predisposes for the serrated CRC pathway, accompanied by common serrated features like gastric metaplasia, a regenerative gene program and classical stem cell depletion^46,47^. Furthermore, elevated expression of our newly generated *LKB1-*mutant gene signature was significantly enriched in SSLs compared to classical adenomas^50^. These findings suggest that PJS cancer pathways differ from the conventional APC-mutant adenocarcinomas in FAP^57^.

A recently presented model proposed that different environmental triggers like microbial dysbiosis and subsequent MAPK pathway activation drive SSL formation^58^. While we currently do not know how external stressors (such as microbial dysbiosis) may affect tumor development in PJS patients, we observed that introduction of oncogenic *Kras* in *Lkb1*-mutant organoids amplifies serrated features and enhances organoid-forming capacity. Furthermore, our results reveal that homozygous *Lkb1* loss induces two organoid phenotypes with either a budding or a cystic morphology accompanied by strongly induced serrated features. Together, our findings suggest that the PJS epithelium is sensitized to events that synergistically drive cells into a premalignant serrated state, such as intestinal damage or activating mutations in MAPK pathway components. Furthermore, as clinical management of PJS patients involves life-long monitoring for polyp resection and (pre)cancerous lesion screening, our results support a rationale for investigating SSL prevalence in PJS cohorts. The malignancy of such lesions may be characterized by markers identified in this study, such as ANXA1.

Together, our study reveals that intestinal organoids are valuable models for *LKB1* mutation effects in PJS patients. Our findings establish a new function of *LKB1*, relating its loss to the induction of a premalignant serrated program which provides a potential explanation for increased cancer incidence in PJS patients. The presented models may provide a starting point to uncover how additional mutations and environmental stressors interact with *LKB1* mutations and how stromal and immune components contribute to malignant growth of the PJS epithelium. Ultimately, in-depth knowledge on the pathways and processes involved in driving PJS epithelia into a pre-malignant state may help to design strategies that re-establish epithelial homeostasis and reduce the incidence of cancer development in PJS patients.

---

**Figure S1.**
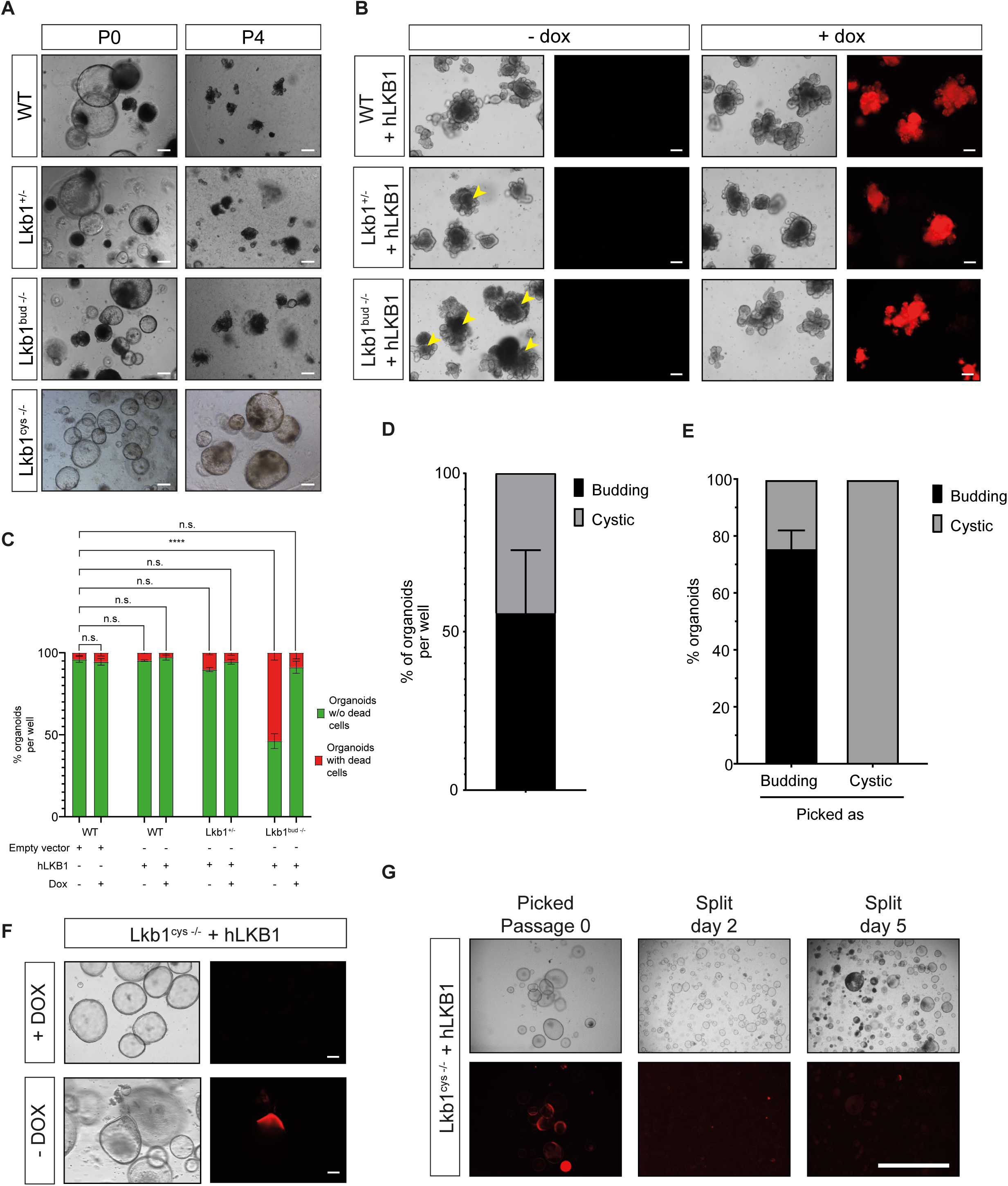

**Figure S2.**
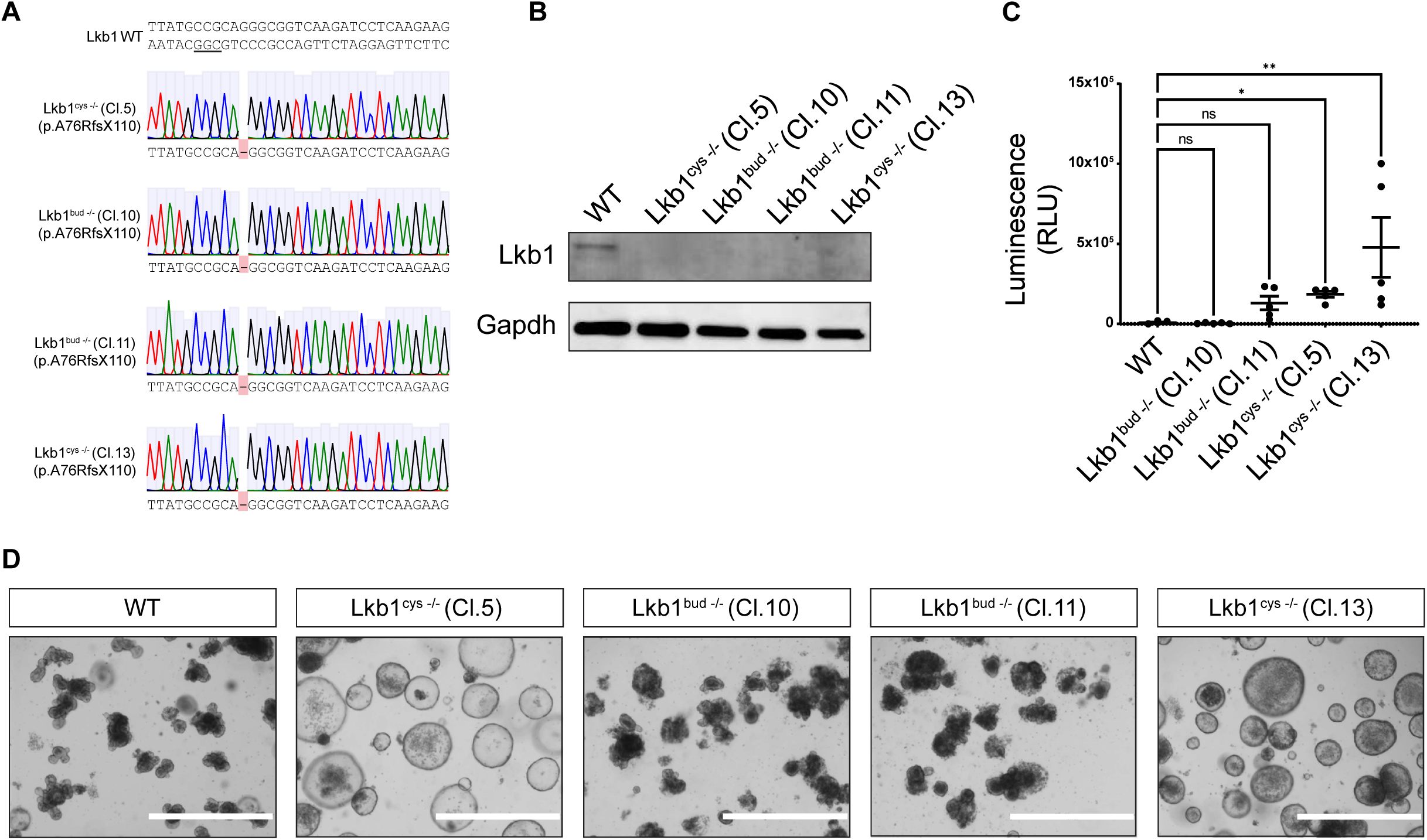

**Figure S3.**
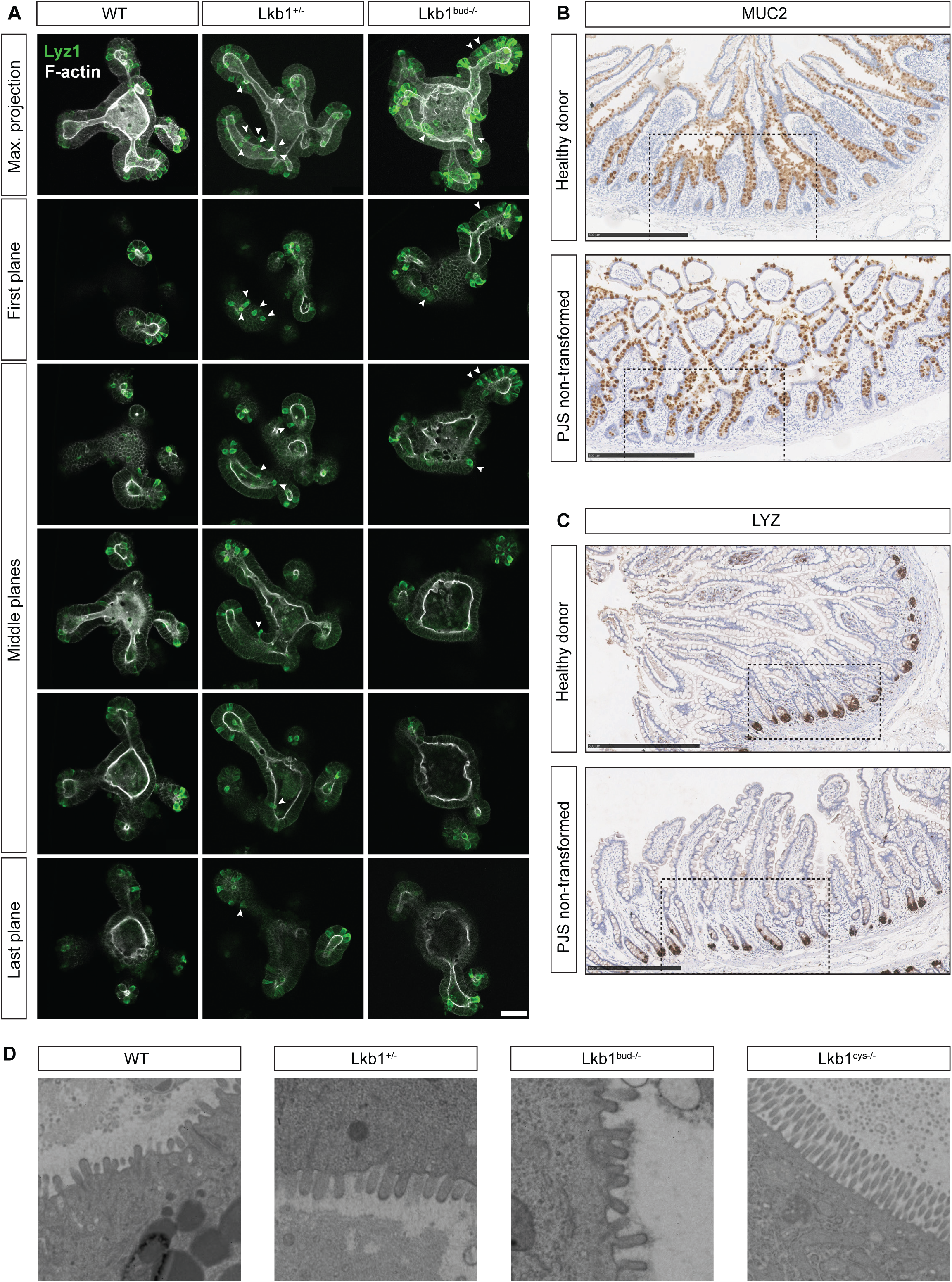

**Figure S4.**
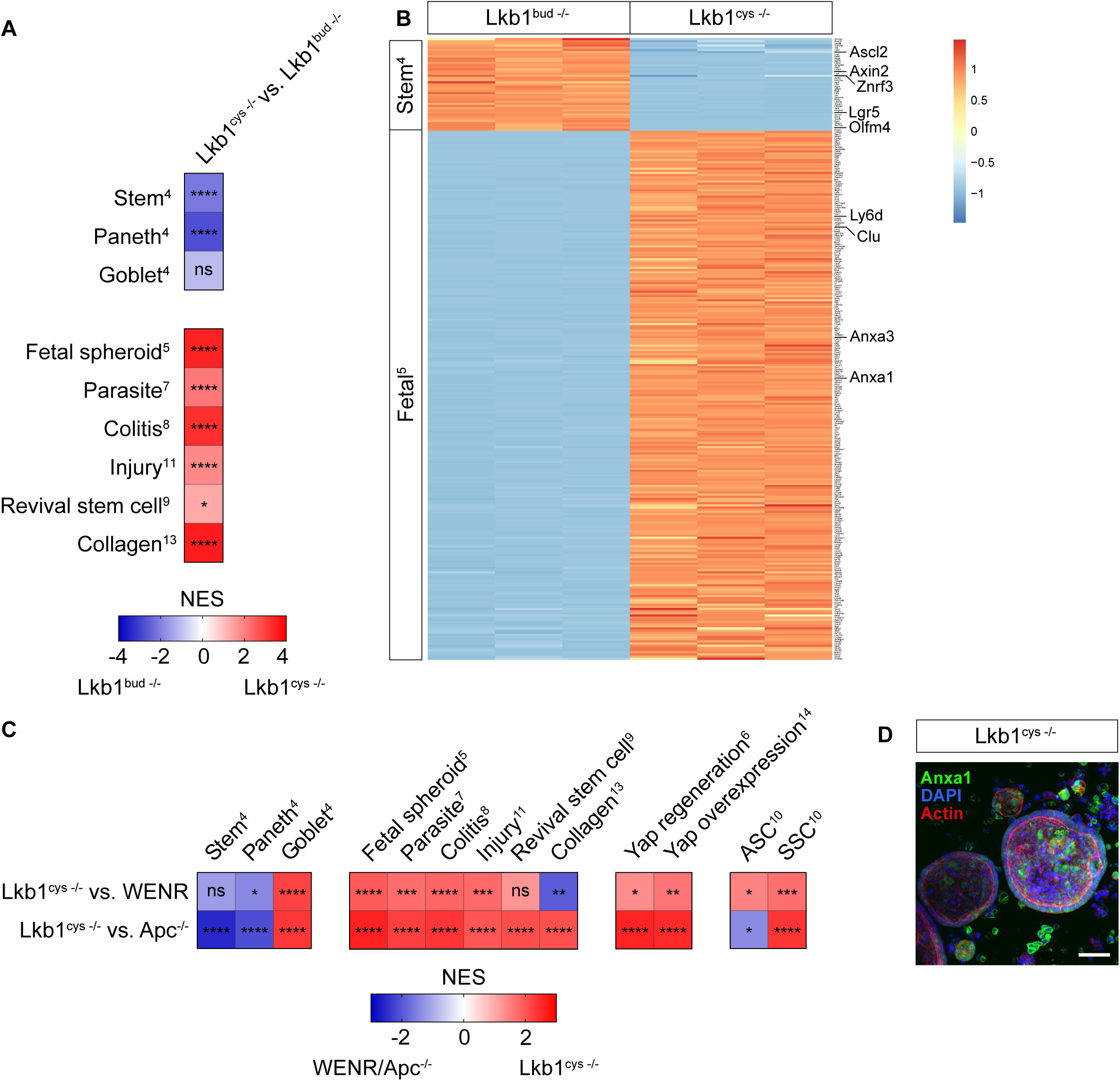

**Figure S5.**
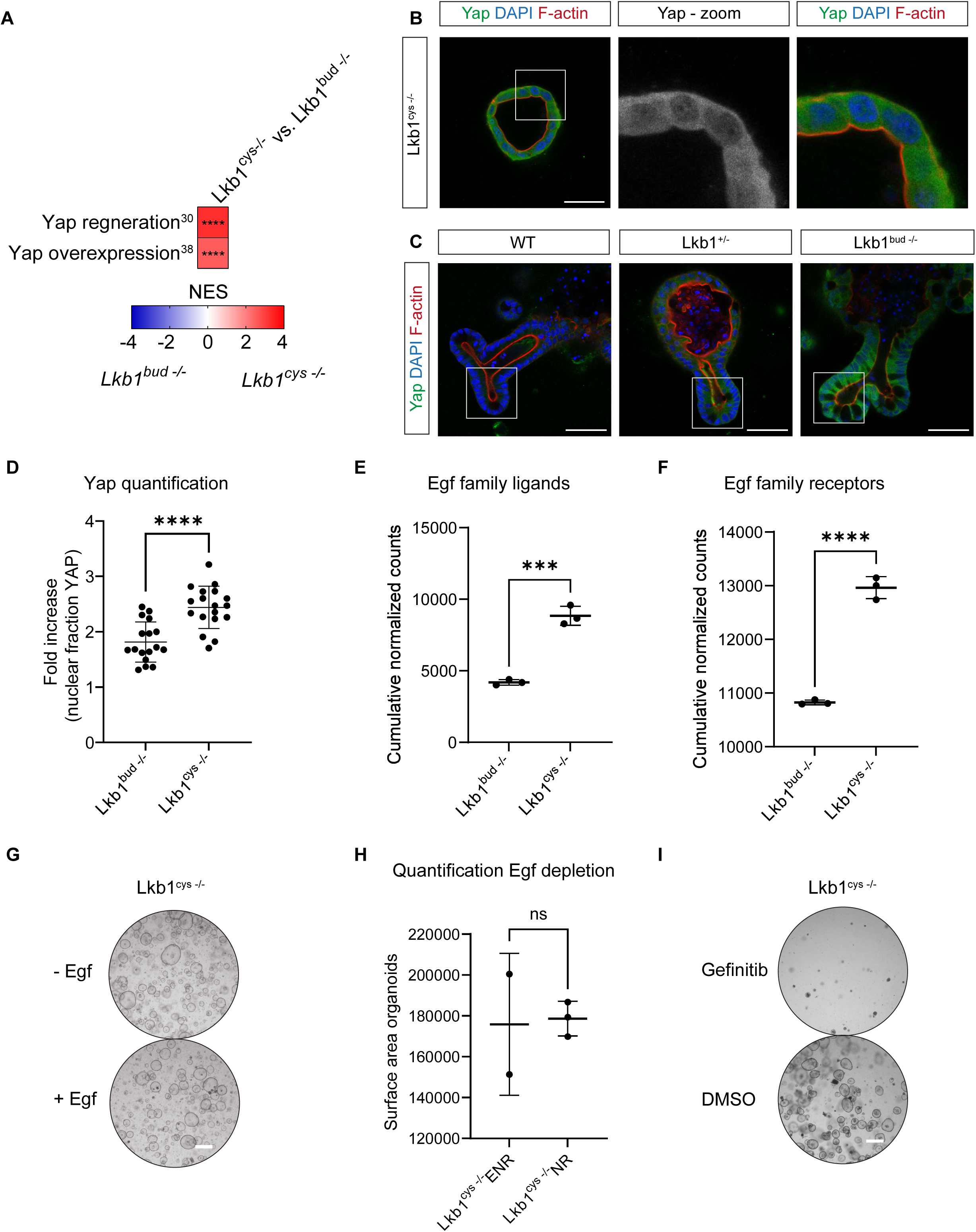

**Figure S6.**
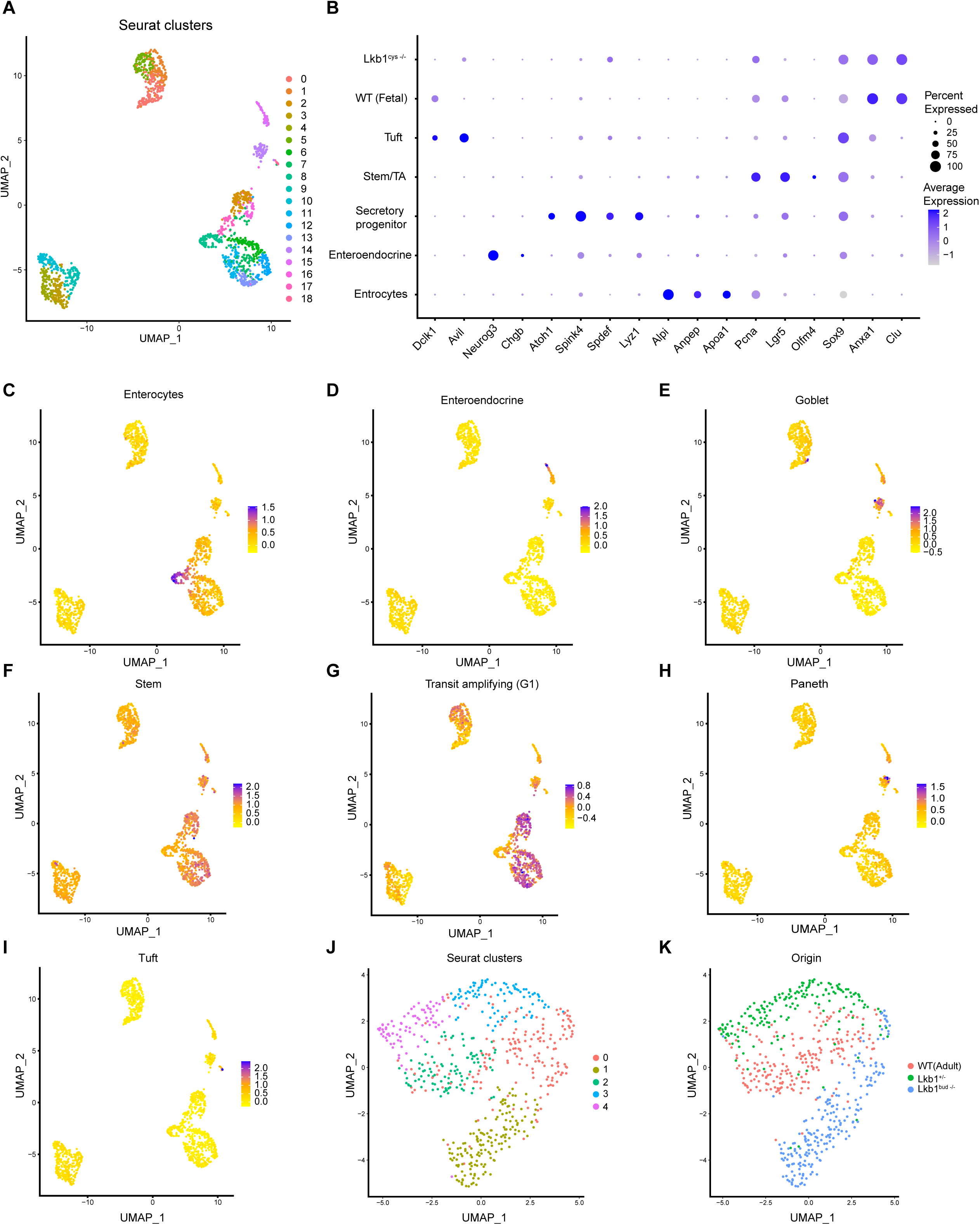

**Figure S7.**
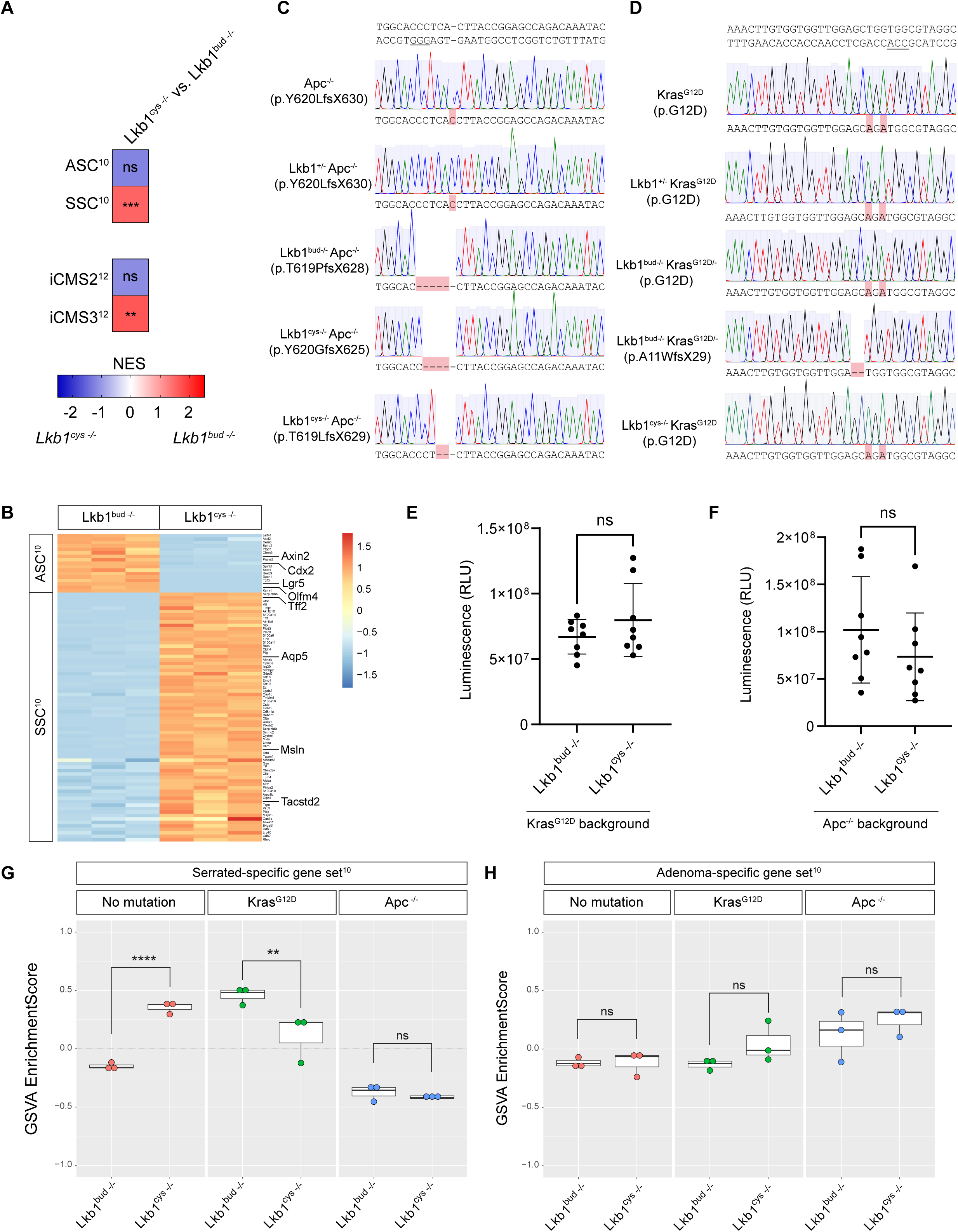

**Figure S8.**
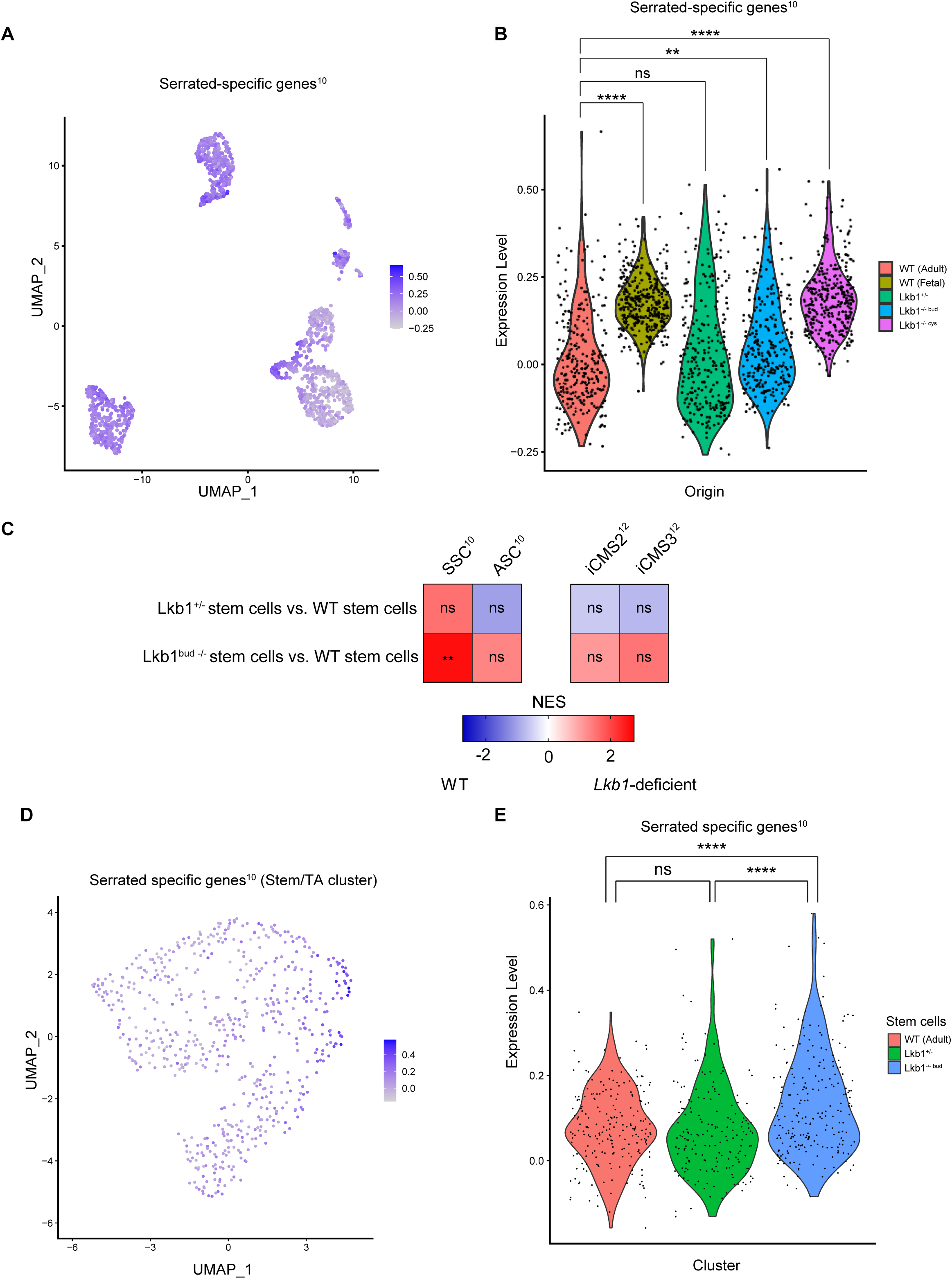

