## Supplementary Materials for "Intestinal LKB1 loss drives a pre-malignant program along the serrated cancer pathway"

**Materials and Methods**

**Patient Material**

In general, patients signed informed consent after ethical committees approved the study protocols. The human colon organoid used can be identified by HUB code HUB-02-A2-040 catalogued at https://huborganoids.nl/ and can be requested at. Distribution to third (academic or commercial) parties will have to be authorized by the Biobank Research Ethics Committee of the University Medical Center Utrecht (TCBio) at request of Hubrecht Organoid Technology (HUB). Formalin-Fixed Paraffin-Embedded (FFPE) intestinal tissue samples from five PJS were obtained from our pathology archives. These samples were collected during routine diagnostic endoscopic procedures. During polyp removal, a separate sample of adjacent non-transformed tissue was also taken. Additionally, five FFPE samples from non-PJS individuals were selected by an experienced pathologist. Tissue sections were stained for MUC2 and LYZ following standard protocols (see “Immunohistochemistry” in supplementary materials). Stained, pseudonymized slides were scanned by the pathology department and transferred for analysis. Quantification was performed as described in the supplementary materials.

**Organoid culture (medium composition)**

Basal culture medium for mouse organoids contained advanced DMEM/F12 (Gibco) supplemented with 1% Penicillin-Streptomycin (Sigma-Aldrich), 10 mM HEPES (Gibco), 1% Ultraglutamine 1 (Lonza), 1x B27 (Sigma-Aldrich), 1x N2 (Sigma-Aldrich) and 1 mM N-acetylcysteine (Sigma- Aldrich). Basal culture medium was further supplemented with 50 ng/mL murine recombinant Egf (Peprotech), 2.5% R-spondin1 (RSPO) conditioned medium (homemade) and 1% Noggin conditioned medium (homemade) to obtain the complete ENR (EGF-Noggin-Rspondin) medium, similar to previous reports^1^. For growth factor withdrawal experiments, EGF was removed from organoid culture medium and organoids were monitored for a total of 16 days. To obtain wild type cystic organoids, wild type organoids were supplemented with 50% Wnt3a-conditioned medium (homemade) and 1mM Nicotinamide (Sigma-Aldrich). Human colon organoids were cultured in advanced DMEM/F12 medium (Invitrogen), supplemented with 1% Penicillin/Streptomycin (P/S, Lonza), 1% Hepes buffer (Invitrogen), 1% Glutamax (Invitrogen), 1x B27 (Invitrogen), 10 mM Nicotinamide (Sigma-Aldrich), 1.25 mM N-acetylcysteine (Sigma-Aldrich), 50 ng/mL EGF (PeproTech), 500 nM TGF-b type I receptor inhibitor A83-01 (Tocris), 10 µM P38 inhibitor SB202190 (Sigma-Aldrich), 0.5% surrogate WNT-CM (homemade), 10% Noggin-CM (homemade), 20% Rspo1-CM (homemade) and 100 µg/mL Primocin (Invitrogen). Bright field images were acquired with the Invitrogen™ EVOS™ FL Color Imaging System (Thermo Fisher Scientific).

**Plasmids**

The pSpCas9(BB)-2A-Puro plasmid was obtained from Addgene (48139). Single guide RNAs (sgRNAs) are listed in **Table S5**. All gRNAs fulfilled the criteria for maximized on target activity and minimized off-target effects^2^. Cloning of sgRNAs into the pSpCas9(BB)-2A-Puro plasmid was performed according to previously published protocols^3^. To generate overexpression cassettes, human LKB1 was subcloned into PiggyBac–CMV–MCS–IRES–mCherry by PCR. All vector sequences were confirmed by Sanger sequencing. To obtain *Lkb1/LKB1*-mutant organoids, a total of 20 µg of DNA (10 µg gRNA, 7.2 µg of pPB-CAG-rtTA_Hygromycin and 2.8 µg PiggyBac transposase) was used. To obtain *Apc* and *Kras* mutant organoids a total of 10 µg DNA (gRNA and gRNA with oligonucleotide respectively) was used.

For generation of overexpression organoids, 5 μg PiggyBac-CMV-hLKB1 overexpression construct, 5 μg rTTA-IRES-Hygro, and 1.7 μg PiggyBac transposase were used to generate organoid lines with inducible overexpression of hLKB1.

**Gene editing of organoids**

Mouse organoids were expanded in ENR medium without antibiotics and supplemented with 50% Wnt3a-conditioned medium (homemade) and 1mM Nicotinamide (Sigma-Aldrich). Human colon organoids were expanded in regular culture medium without antibiotics. Organoids were electroporated using a NEPA21 (NEPAGENE) electroporator. 100 µg/mL Hygromycin B was used to select *Lkb1/LKB1*-mutant organoids. After selection, single organoids were manually picked for clonal expansion and then genotyped. Genomic DNA was isolated using the DNA micro kit (Qiagen) according to the manufacturer’s instructions and PCR amplification was performed using the GoTaq Flexi DNA polymerase kit (Promega). Organoid genotype was confirmed by Sanger sequencing. Primers are listed in **Table S6.**

For selection of *Apc* and *Kras* mutant organoids, RSPO and Egf were omitted from the culture medium respectively.

**Morphology quantification**

After obtaining a 100% knockout-score, as determined by Synthego tool, using sgRNA3, organoids were cultured for 4 weeks in ENR. Organoids were mechanically sheared and plated in minimal density to optimize quantification. Six individual wells were imaged using Invitrogen™ EVOS™ FL Color Imaging System (Thermo Fisher Scientific). Number of budding and cystic organoids was manually determined for each well on day 5 after splitting. Each well contained between 9 and 31 organoids (median = 21). From these bulk cultures, nine budding organoids and nine cystic organoids were manually picked and expanded as clonal lines for two passages (two weeks). These cultures were imaged using Invitrogen™ EVOS™ FL Color Imaging System (Thermo Fisher Scientific) and once again manually counted based on morphology on day 5 after splitting. Each well contained between 13 and 41 organoids (median = 18).

**Analysis of overexpression organoid lines**

1 μg/ml doxycycline was added to induce overexpression of hLKB1 in the clonal lines. Organoids were mechanically split using standard procedures and treated with 1 μg/ml doxycycline for 14 days. After 5 and 10 days of treatment, the organoids were mechanically split using standard procedures. Brightfield images were taken on day 4 of split 2 (day 14). Quantification of the organoid phenotypes was performed on brightfield images manually.

**Protein extraction and Western blotting**

Organoids were harvested three days after splitting and washed in cold PBS and collected in Cell Recovery Solution (Corning) according to the manufacturer’s instructions. Whole cell lysates were obtained by direct lysis in RIPA buffer (50 mM Tris- HCl pH 8.0, 150 mM NaCl, 0.1% SDS, 0.5% Na-Deoxycholate, 1% NP-40) containing complete protease inhibitors (Sigma-Aldrich) for 30 minutes on ice. Proteins were run on SDS–PAGE gels and transferred to PVDF membranes, followed by blocking with Odyssey blocking buffer (LI-COR). The antibodies used are described in **Table S7**. Blots were imaged with the Amersham Typhoon (GE Healthcare Life Sciences).

**Immunofluorescence**

Organoids were seeded in 15-well Ibidi chambers after passaging and fixed 3-4 days later in 4% Paraformaldehyde (Sigma-Aldrich). After fixation organoids were washed with cold PBS and permeabilized with PBS containing 0.2% Triton, 1% DMSO and 1% BSA (PBD0.2T buffer) and incubated with primary antibodies diluted in PBD0.2T buffer at 4°C overnight. Washes were performed with PBD0.2T buffer and secondary antibodies were incubated 2-3 hours at room temperature. The antibodies used are described in **Table S7**. Images were acquired with a Zeiss LSM700 or LSM880 confocal microscope.

**Yap quantification**

Quantification of the nuclear fraction of Yap was analyzed using the JACoP Fiji plugin using DAPI as nuclear marker.

**Electron microscopy**

Three days after splitting, organoids were fixed in a solution of 2% paraformaldehyde and 2.5% glutaraldehyde in 0.1 M phosphate buffer (pH 7.4) overnight at 4°C. On the following day, organoids were washed twice in 0.1 M phosphate buffer (pH 7.4) and subsequently post-fixed with 1% Osmium tetroxide for 2 hours at 4°C. Organoids were then stained with 0.5% uranyl acetate 1 hour at 4°C in the dark. Matrigel droplets were transferred into glass vials for dehydration with increasing acetone concentrations according to the following protocol: 70% overnight at 4°C, 90% for 15 minutes at RT, 96% for 15 minutes at RT, and 100% 3 times for 30 minutes each at RT. After dehydration, samples were embedded in EPON resin. Ultrathin sections of 65 nm were obtained with a Leica UC7 microtome and stained with uranyl and lead citrate. Electron microscopy was performed in the Cell Microscopy Core (CMC) of the UMC Utrecht. Sections were imaged with a JEOL JEM-1010 transmission electron microscope (TEM).

**Immunohistochemistry**

Fresh mouse small intestinal organoids were fixed in 4% formaldehyde three days after splitting for 1 hour followed by dehydration in 25%, 50% and 70% ethanol and embedding in paraffin. 4μm FFPE organoid or PJS patient tissue sections were obtained, deparaffinized, blocked in 0.3% H2O2 in methanol for 20 minutes. Antigen retrieval was performed in Tris/EDTA buffer (10 mM/1 mM; pH 9.0) or sodium-citrate buffer (0,01 M/pH6.0) for 20 minutes at 100°C. Nonspecific binding sites were blocked using Protein Block Serum-free (DAKO) for 10 minutes, followed by primary antibody incubation. Antibody binding was visualized using the using BrightVision Goat-anti-Rabbit/Poly-AP and BrightVision Goat-anti-Mouse/Poly-HRP (VWR/ImmunoLogic), with DAB as chromogen (Sigma-Aldrich). Sections were counterstained with Haematoxylin or neutral red and mounted using Pertex. Primary antibodies used are described in **Table S7.**

**Counting patient tissue samples**

In 10 microscopically normal looking crypts MUC2^+^ and LYZ^+^ cells were counted from the bottom toward to the top of the crypt. Only longitudinally oriented crypts that were completely visible on the slide were counted. Cell positions were defined as the bottom of the crypt is cell position one (1), going upward to cell position +2, +3, and so forth till cell position 30. Within the part of the crypt between cell position 1 and 30, the number of positively labelled cells were counted. Also, the total cell position was determined by adding up each cell position of a labelled cell and average cell position of the labelled cells was determined by adding up each cell position of a labelled cell and dividing the sum by the total number of labelled cells counted. For MUC2+ cell quantification, 40 crypts from 4 individuals of healthy donors and 30 crypts from 3 PJS individuals were counted. For LYZ+ cell quantification, 50 crypts from 5 individuals of healthy donors and 30 crypts from 3 PJS individuals were counted.

**Cell viability assay**

For the organoid reconstitution assays, mouse small intestinal organoids were dissociated to single cells after incubation with TrypLE (Life Technologies) for 5 min at 37°C. Dissociated organoids were passed through a 40 μm strainer and counted. Cells were seeded at a density of 5000 cells/10μl Matrigel and cultured in ENR medium with 10 μM Y-27632 (Sigma-Aldrich) for 7 days.

For EGF withdrawal experiments, human colon organoids were dissociated to single cells and filtered similarly to mouse small intestinal organoids. Cells were seeded at a density of 1000 cells/μl in 25μl BME droplets. Organoids were then lysed with CellTiter-Glo® (Promega) according to the manufacturer’s protocol. ATP produced by viable cells was measured by a Berthold luminometer Centro LB960.

**Bulk RNA sequencing**

Organoids were lysed in RLT lysis buffer (Qiagen) 3-5 days after passaging. Biological triplicates were employed. To isolate RNA from samples, the “miRNA CT 400” protocol was run on a QIAsymphony isolation robot using the QIAsymphony RNA Kit (931636) on each sample in 400ul of the RLT Plus lysis buffer (1053393). RNA quality was checked with the Agilent Fragment Analyzer 5300 system using the RNA Kit (15nt) (Cat. DNF-471-1000). And RNA quantity was measured with the Invitrogen™ Qubit ™ Fluorometer using the Qubit RNA HS Assay Kit (Cat. Q32855). 100ng of total RNA was used to prepare TruSeq Stranded mRNA libraries (Cat. 20020594) following the manufacturers protocol, with full (WENR, WT, Lkb1^+/-^, Lkb1^bud-/-^ and Lkb1^cys-/-^ organoids) or half (Lkb1^mut^ + Apc^-/-^ and Lkb1^mut^ + Kras^G12D^ organoids) the reagents volume.

After the library preparation libraries were checked with the Fragment Analyzer system dsDNA 910 Reagent Kit (35-1500bp) (Cat. DNF-910-K1000) and with Qubit dsDNA HS Assay Kit (Cat. Q32854). Sample libraries were pooled equimolar. Samples were sequenced on Ilumina NextSeq500 (1x 75bp length) (WENR, WT, Lkb1^+/-^, Lkb1^bud-/-^ and Lkb1^cys-/-^ organoids) or Ilumina NextSeq2000 (2x 50bp length) (Lkb1^mut^ + Apc^-/-^ and Lkb1^mut^ + Kras^G12D^ organoids).

RNA isolation and library prep of human colon organoids was performed similarly to mouse organoids using half volumes of the reagents. Samples were sequenced on Ilumina NextSeq2000 (2x 50bp length).

**Bulk RNA sequencing data analysis**

Single-end RNASeq reads were processed using the UMCU RNASeq pipeline (<https://github.com/UMCUGenetics/RNASeq>, v2.3.0) with default settings. Read quality was assessed by FastQC (0.11.4) followed by splice-aware alignment against the mouse reference genome (GRCm38) or human genome (GRCh38) with STAR (2.4.2a). RNA expression quantification was performed with htseq-count (0.6.0) in reverse-stranded mode. Multiple sequence runs were integrated using standardized integration parameters. Differential gene expression analysis was performed with the DESeq2 (1.22.2) R package. In each comparison, genes were selected with an absolute fold-change > 2 and adjusted p-value < 0.05. Data clustering and generation of heatmaps was performed with pheatmap. Gene Set Enrichment Analysis was performed using GSEA (v4.2.0, Windows App) and visualized using ggplot2. Intestinal cell-type specific gene sets for GSEA were obtained from Extended Figure Table 3 of Haber et al., 2017^4^. The other datasets employed in GSEA were derived from previously published studies^5,6,15,7–14^. Human gene sets were converted to their respective mouse homologs using biomaRt (2.50.3) in R where necessary.

**Single cell RNA sequencing**

After single cell dissociation using TrypLE at 37°C cells were filtered through a 40 µm nylon cell strainer (Falcon). Variable single cells were FACS sorted into 384-well cell capture plated (Single Cell Discoveries). Each well of a cell capture plate contains a small 50nl droplet of barcoded primers and 10µl of mineral (Signa, M8410). After sorting, plates were immediately spun and placed on dry ice. Plated were stored at -80ºC.

scRNA-seq was performed by Single Cell Discoveries according to an adapted version of the SORT-seq protocol^16^ with primers described in earlier^17^. Cells were heat-lysed at 65ºC followed by cDNA synthesis. After second-strand cDNA synthesis, all the barcoded material from one plate was pooled into one library and amplified using *in vitro* transcription (IVT). Following amplification, library preparation was done following the CEL-Seq2 protocol^18^ to prepare a cDNA library for sequencing using TruSeq small RNA primers (Illumina). The DNA library was paired-end sequenced on an Illumina NextSeq500 at paired-end 60- and 26-bp read length and 75,000 reads per cell.

**Single cell RNA sequencing analysis**

*Mapping and filtering*: The Sharq pipeline was used to process the sequencing data^19^. Mapping was performed using STAR (version 2.6.1), on the Genome Reference Consortium Mouse built 38 (GRCm38). Read assignment was with feature Counts version 1.5.2 using a gene annotation based on GENCODE version M14. External RNA controls (ERCCS) and transcripts mapping to the mitochondrial genome were removed from all cells. Low quality cell barcodes having either less than 800 unique transcripts, 200 genes 33.3% mitochondrial transcripts or over 4-fold non-exonic to exonic reads were excluded, as well as barcodes with over 100,000 unique transcripts.

*scRNA-seq analysis*: Unique transcript counts were normalized using sctransform^20^ and analyzed using the Seurat R package (version 4.1.0)^21^. From the top 3000 variable genes, genes associated with cell cycle phase, dissociation stress (heat shock and chaperone proteins according to GO:0006986), sex (Xist, Tsix, and Y chromosome-specific genes), and activity (ribosomal protein genes according to GO:0022626) were removed from the list of variable genes to avoid biases in cell clustering, as described before^22^. The first 20 principal components were used both to calculate dimensionality reduction using UMAP and to perform clustering with a resolution of 1, k.param = 10 and the Louvain algorithm.

**TCGA data analysis**

Signature scores for a *LKB1*-mutant geneset were calculated for each sample as log2 Z-score over the samples from the TCGA Colon Adenocarcinoma (2022 – v32) set or the Tumor Colon (tubular-serrated) (GSE45270^23^) set using the R2 genomic analysis platform (<http://r2.amc.nl>). *LKB1*-mutant geneset was determined using gene dosage of 0 (WT), 1 (*LKB1^+/-^*) and 2 (*LKB1^-/-^*) as design input in DESeq2 (version 1.48.0) R package. *LKB1*-mutant geneset contains all genes that were significantly upregulated (padj<0.05).

**Table S5: gRNA sequences**

| Gene | Sequence |
| --- | --- |
| *Lkb1* gRNA1 | AGCTTGGCGCGTTTGCGGCG |
| *Lkb1* gRNA2 | TGAGGATCTTGACCGCCCTG |
| *Lkb1* gRNA3 | CATCGGCAAGTACCTGATGG |
| *LKB1* | GTTGCGAAGGATCCCCAACG |
| *Apc* | TGTCTGGCTCCGGTAAGTGA |
| *Kras* | CTTGTGGTGGTTGGAGCTGG |

**Table S6: Genotyping primers**

| Gene | Forward primer | Reverse Primer |
| --- | --- | --- |
| *Lkb1*  gRNA1/3 | GGGGAAAATCAAAAGTGAAGAA | CCCTCTAGCTGCGTAAACAAAC |
| *Lkb1*  gRNA2 | CTCCACCGAGGTAATCTACCAGCCG | TCTAACCGTTTTCTCTTCTCATTTCC |
| *LKB1* | ATCGACTCCACCGAGGTCATCTAC | CCAGCTCAGGGTGTTAAGAGGAAGT |
| *Apc* | ATAGAAACAGCACTGACCCAAATTTCA | AGGCCTCTTTGCTTAGAGCTTTCATAA |
| *Kras* | TGGCTCCAACACAGATGTTC | GGATGGCATCTTGGACCTTA |

**Table S7: Western Blot antibodies**

| Antibody | Dilution | Supplier | Cat. number |
| --- | --- | --- | --- |
| Anti-LKB1 | 1:1000 | Santa Cruz | sc-32245 |
| Anti-GAPDH | 1:8000 | Millipore | CB1001 |
| Anti-mouse Alexa 800 | 1:5000 | LI-COR | 926-32211 |
| Anti-rabbit Alexa 680 | 1:5000 | LI-COR | 925-68070 |
| Anti-pAMPK | 1:1000 | Cell Signaling | 2535 |
| Anti-AMPK | 1:1000 | Cell Signaling | 2793 |
| Anti-Actin | 1:10000 | MP Biomedicals | 691001 |
| Anti-4EBP1 | 1:1000 | Cell Signaling | 9452 |
| Anti-Vinculin | 1:5000 | Abcam | ab129002 |

**Table S8: Immunostaining antibodies**

| Antibody | Dilution | Supplier | Cat. number |
| --- | --- | --- | --- |
| Anti-Muc2 | 1:200 (IF)  1:1000 (IHC) | Santa Cruz | sc-515032 |
| Anti-Lysozyme1 | 1:200 (IF)  1:2000 (IHC) | Dako | A0099 |
| Anti-ANXA1 | 1:200 (IF) | Atlas Antibodies | HPA011271 |
| Anti-YAP | 1:200 (IF)  1:200 (IF) | Santa Cruz  Millipore | sc-101199  MABS2029 |
| Anti-Cleasved-Caspase-3 | 1:200 (IF) | Cell Signalling | 9661 |
| Anti-rabbit Alexa Fluor 488 | 1:300 (IF) | Invitrogen | A11034 |
| Anti-mouse Alexa 488 | 1:300 (IF) | Invitrogen | A11029 |
| Anti-rat Alexa 488 | 1:300 (IF) | Invitrogen | A11006 |
| DAPI | 1:1000 (IF) | Sigma | D9542 |
| Click-iT™ EdU Alexa Fluor™ 647 | n.a. | Invitrogen | C10424 |
| [Alexa Fluor™ 647 Phalloidin](https://www.thermofisher.com/order/catalog/product/A22287?SID=srch-srp-A22287) | 1:500 (IF) | Invitrogen | A22287 |

**Acknowledgements**

The authors thank members of the Maurice laboratory for helpful discussions and suggestions. PiggyBac Transposase and pPB-CAG-rtTA_Hygromycin were a kind gift from B.K. Koo. Graphical abstract was created in BioRender. Maurice, M. (2025) <https://BioRender.com/5tt6n2a>. We acknowledge the Cell Microscopy Core (CMC) of the Center for Molecular Medicine, UMC Utrecht for providing microscopy training and service. We acknowledge the Utrecht Sequencing Facility (USEQ) for providing sequencing service and data. USEQ is subsidized by the University Medical Center Utrecht and The Netherlands X-omics Initiative (NWO project 184.034.019).

**Figure S1. Phenotypic reversal upon LKB1 WT overexpression.**

**A)** Representative brightfield images of different clonal organoid lines after electroporation followed over time in ENR medium. In passage 0 (start of ENR culture; P0) all lines show cystic morphology. After four passages in ENR (P4), most clonal lines reverted into budding organoids expect for some *Lkb1^-/-^* organoids. **B)** Brightfield and fluorescent images of mCherry-hLKB1 organoids in WT, *Lkb1^+/-^* and *Lkb1^bud-/-^*organoids. Expression of hLKB1 after doxycycline induction is visualized by mCherry expression. Yellow arrowheads indicate organoids with apoptotic cells surrounding the organoids. This phenotype is lost after induction of hLKB1. Scale bar = 100µm. **C)** Bar plot indicating the percentage of organoids without (green) and with (red) apoptotic cells per well as identified in brightfield microscopy. *N* = 5 (WT), *N* = 8 (Lkb1^+/-^) and *N* = 10 (Lkb1^bud-/-^). Bar plots show mean value ± SEM. ns = not significant, **** = p≤0.0001. **D)** Bar plot indicating the percentage of organoids in cystic or budding morphologies in bulk cultures (100% knock-out score (gRNA3)) (*N* = 6). Bar shows mean ± SD. **E)** Bar plot indicating the percentage of organoids in cystic or budding morphologies in clonal cultures picked as either budding (*N* = 9) or cystic (*N* = 9). Bar shows mean ± SD. **F)** Brightfield and fluorescent images of mCherry-hLKB1 organoids in *Lkb1^cys-/-^* organoids. Expression of hLKB1 after doxycycline induction is visualized by mCherry expression. Scale bar = 100µm. **G)** Representative images of doxycycline-treated Lkb1^cys-/-^ organoids to induce hLKB1 overexpression (+OE). mCherry^+^ organoids were manually picked and plated (left panel). mCherry expression was diminished in organoids within 2 days after splitting (middle panel) and completely lost 5 days after splitting (right panel). Scale bar = 1mm.

**Figure S2. *Lkb1* loss induced by different gRNAs results in similar heterogeneous phenotypes.**

**A)** Sequences of *Lkb1*-mutant organoids obtained with gRNA2 showing indels/deletions detected in exon 1 of the *Lkb1* gene. PAM sequences are underlined. **B)** Western blot analysis of Lkb1 expression in WT and indicated *Lkb1*-mutant organoids. **C)** Organoid reconstitution assay from single cells for wild type (WT) and indicated Lkb1^-/-^ organoids; cell viability was measured at day 7. Friedman test was applied for analysis. ns = not significant, *** = *p < 0.05*, ** = *p < 0.01*. Each dot represents a biological replicate, *N* = 5. **D)** Representative brightfield images of WT and indicated *Lkb1*-mutant organoids are shown. Scale bar = 1mm.

**Figure S3. Goblet and Paneth cell number and positioning in *Lkb1*-mutant organoids and PJS epithelia.
A)** Immunofluorescence of Lysozyme1 (Lyz1, green) in WT and indicated *Lkb1*-mutant clones. Cellular membranes are visualized using F-actin (white). Images are max projections of z-stack (top panel) and single plane images. White arrows indicate mislocalized Lyz1^+^ cells. Cellular membranes are visualized by F-actin (white). Multiple planes of a single z-stack are shown. Images of are digitally enhanced for optimal visualization. Scale bar = 50 μm. **B-C)** Overview images of immunohistochemistry staining of MUC2 **(B)** and LYZ **(C)** in healthy and PJS non-transformed epithelium. Zoom-in images in Figure 2 are indicated with dotted boxes. Scale bar = 500 µm. **D)** Zoomed in electron microscopy images to show apical brush border formation in wild type (WT) and indicated *Lkb1*-mutant organoids.

**Figure S4. *Lkb1^cys-/-^* organoids display increased expression of a regenerative program in comparison to *Lkb1^bud-/-^* organoids.**

**A)** *Lkb1*^cys-/-^ versus *Lkb1^bud-/-^* organoid derived bulk mRNA-seq sequencing data were analyzed by gene set enrichment analysis (GSEA) for cell types^4^, colitis^8^, injury^11^, parasite^7^, fetal spheroid^5^, collagen^13^ and revival stem cell^9^ gene sets. Heatmap displays normalized enrichment scores (NES). ns = not significant, * = FDR < 0.05, *** = FDR ≤ 0.001, **** = FDR ≤ 0.0001. **B)** Heatmap of z-score transformed expression values of *Lgr5^+^* stem cell genes (Stem)^4^ enriched in *Lkb1^bud-/-^* organoids (top cluster) and fetal genes (Fetal)^5^ enriched in *Lkb1^cys-/-^* organoids (bottom cluster). Key marker genes are highlighted. Each column represents a biological replicate of the indicated group, *N* = 3. **C)** *Lkb1*^cys-/-^ versus *Apc*^-/-^ or Wnt-treated (WENR) organoid derived bulk mRNA-seq sequencing data were analyzed by gene set enrichment analysis (GSEA) for cell types^4^, colitis^8^, injury^11^, parasite^7^, fetal spheroid^5^, collagen^13^, revival stem cell^9^, Yap^6,14^, adenoma specific^10^ (ASC) and serrated specific^10^ (SSC) gene sets. Heatmap displays normalized enrichment scores (NES). ns = not significant, * = FDR < 0.05, ** = FDR ≤ 0.01, *** = FDR ≤ 0.001, **** = FDR ≤ 0.0001. **D)** Immunofluorescence staining of the fetal marker Anxa1 (green) in *Lkb1*^cys-/-^ organoids. Cellular membranes are visualized by F-actin (red) and nuclei by DAPI (blue). Scalebar = 50 μm**.**

**Figure S5. *Yap* and *Egf* expression are enhanced in *Lkb1^cys-/-^* organoids compared to *Lkb1^bud-/-^* organoids.**

**A)** *Lkb1*^cys-/-^ versus *Lkb1^bud-/-^* organoid-derived bulk mRNA-seq sequencing data were analyzed by gene set enrichment analysis (GSEA) for Yap gene sets during regeneration^6^ or overexpression^14^. Heatmap displays normalized enrichment scores (NES). ns = not significant, * = FDR < 0.05, *** = FDR ≤ 0.001, **** = FDR ≤ 0.0001. **B)** Immunofluorescence staining of Yap (green) in *Lkb1^cys-/-^* organoids. Nuclei are visualized by DAPI staining (blue) and cellular membranes are visualized by F-actin (red). High magnification of the area indicated by white box is shown in the lower two panels. Scale bar = 50 µm. Representative images of N = 6 experiments are shown. **C)** Zoomed out immunofluorescence staining for Yap (green), DAPI (blue) and F-actin (red) in WT, *Lkb1*^+/-^ and Lkb1^bud-/-^organoids. White box corresponds to the area shown in Figure 4B. Scale bar = 50 µm. **D)** Quantification of the nuclear fraction of Yap. Dots represent individual organoids derived from four independent experiments. *N* = 17 for *Lkb1*^bud-/-^ organoids and *N* = 18 for *Lkb1*^cys-/-^ organoids. An unpaired t-test was applied for analysis and statistical significance was defined as ***** = p < 0.0001.* **E-F)** Cumulative normalized counts for *Egfr* binding ligands **(E)** and *Egf* family receptors **(F)** in *Lkb1^bud-/-^* and *Lkb1^cys-/-^* organoids. An unpaired t-test was applied for analysis and statistical significance was defined as **** = p < 0.001 and **** = p < 0.0001.* Each dot represents one biological replicate, N = 3. **G)** Representative bright field images of *Lkb1^cys-/-^* organoids grown in the presence (ENR) and absence (NR) of Egf. Organoids were incubated for 7 days in indicated medium and split on day 4. Scale bar = 500 µm. **H)** Calculation of the surface area covered by *Lkb1^cys-/-^* organoids grown in the presence (ENR) and absence (NR) of Egf using the OrganoSeg tool^24^. Each dot represents one biological replicate, N = 2-3. An unpaired t-test was applied for analysis and statistical significance was defined as *ns = not significant.* **I)** Representative bright field images of *Lkb1^cys-/-^* organoids cultured in NR and treated with DMSO or 100 nM Gefitinib. Organoids were incubated for 9 days in indicated medium and split on day 6. Scale bar = 500 µm.

**Figure S6. Identification of intestinal cell types in WT and *Lkb1*-mutant organoids.**

**A)** UMAP visualization of scRNA-seq data from wild-type (WT), fetal and indicated *Lkb1*-mutant organoids. Colors indicate different clusters as defined by Seurat analysis **B)** Expression of known markers for different cell populations in WT, *Lkb1*^+/-^, *Lkb1*^bud-/-^ and *Lkb1*^cys-/-^ organoids. Size of the dot represents percentage of cells expressing marker gene and the color the average expression within that population. **C-I)** Module scores for enterocyte **(C)**, enteroendocrine **(D)**, goblet **(E)**, stem **(F)**, transit amplifying (G1) **(G)**, Paneth **(H)** and tuft **(I)** cell gene signatures^4^ overlaid on the UMAP plot of *Lkb1-*mutant, fetal and WT organoids. **J-K)** Reclustering of stem cell populations from WT, *Lkb1*^+/-^, *Lkb1*^bud-/-^ organoids showing clusters as identified by Seurat analysis **(J)** and by organoid line **(K)**.

**Figure S7. Serrated gene set expression is enhanced in *Lkb1^cys-/-^* organoids compared to *Lkb1^bud-/-^* organoids.**

**A)** *Lkb1^cys-/-^* versus *Lkb1^bud-/-^* organoid derived bulk mRNA-seq sequencing data was analyzed by gene set enrichment analysis (GSEA) for serrated-specific^10^, adenoma specific^10^, iCMS2^12^ and iCMS3^12^ genesets. Heatmap displays normalized enrichment scores (NES). ns = not significant, * = FDR < 0.05, ** = FDR ≤ 0.01, *** = FDR ≤ 0.001, **** = FDR ≤ 0.0001. **B)** Heatmap of z-score transformed expression values of adenoma-specific genes (ASC)^10^ enriched in *Lkb1^bud-/-^* organoids (top cluster) and serrated-specific genes (SSC)^10^ enriched in *Lkb1^cys-/-^* organoids (bottom cluster). Each column represents a biological replicate of the indicated group, N = 3. **C)** Sequences of *Apc-*mutant, *Apc/Lkb1*-mutant, *Kras*-mutant or *Kras/Lkb1*-mutant organoids showing indels/deletions detected in exon 15 of the *Apc* gene or deletions/base substitutions in exon 2 of the *Kras* gene. PAM sequences are underlined. **E)** Organoid reconstitution assay from single cells (analyzed at day 7) shown for *Apc^-/-^/Lkb1^bud-/-^* and *Apc^-/-^/Lkb1^cys-/-^* organoids. Each dot represents a biological replicate, *N* = 8. Unpaired t-test was applied for analysis. *ns = not significant*. **F)** Organoid reconstitution assay from single cells (analyzed at day 7) for *Kras^G12D^/Lkb1^bud-/-^* and *Kras^G12D^/Lkb1^cys-/-^* organoids. Each dot represents a biological replicate, *N* = 8. Unpaired t-test was applied for analysis. *ns = not significant*. **G-H)** Gene Set Variation Analysis (GSVA) analysis of SSC^10^ genes **(G)** and ASC^10^ genes **(H)** for *Lkb1^bud-/-^, Lkb1^cys-/-^, Kras^G12D^/Lkb1^bud-/-^, Kras^G12D^/Lkb1^cys-/-^, Apc^-/-^/Lkb1^bud-/-^* and *Apc^-/-^/Lkb1^cys-/-^* organoids. Each dot represents a biological replicate of the indicated group, N = 3. *ns = not significant, * = p < 0.05, ** = p < 0.01 and **** = p < 0.0001* using one-way ANOVA with Wilcoxon test.

**Figure S8. Induction of serrated genes in *Lkb1*-mutant organoids is reflected at the single cell level. A)** Module score for a serrated-specific (SSC) gene set^10^ on the UMAP plot of indicated *Lkb1*-mutant, fetal and WT organoids. **B)** Violin plot showing a module score for a serrated-specific gene set across WT, fetal, *Lkb1^+^*^/-^, *Lkb1^bud-/-^* and *Lkb1^cys-/-^* organoids. ns = not significant, ** = p < 0.01 and **** = p < 0.0001 using one-way ANOVA with Wilcoxon test. **C)** GSEA was performed for adenoma- and serrated-specific gene set^10^ and CMS-subtypes^12^. Heatmap displays normalized enrichment scores (NES). ns = not significant, ** = FDR ≤ 0.01. **D)** Module score for a serrated-specific gene set^10^ on the UMAP plot of WT stem cells, *Lkb1^+^*^/-^ stem cells, *Lkb1^bud-/-^* stem cells and all *Lkb1^cys-/-^* cells. **E)** Violin plot showing module scores of a serrated-specific gene set^10^ for WT stem cells, *Lkb1^+^*^/-^ stem cells, *Lkb1^bud-/-^* stem cells and all *Lkb1^cys-/-^* cells. ns = not significant, **** = p ≤ 0.0001 using one-way ANOVA with Wilcoxon test.
